# Evidence that variation in root anatomy contributes to local adaptation in Mexican native maize

**DOI:** 10.1101/2023.11.14.567017

**Authors:** Chloee M. McLaughlin, Meng Li, Melanie Perryman, Adrien Heymans, Hannah Schneider, Jesse R. Lasky, Ruairidh J. H. Sawers

## Abstract

Mexican native maize (*Zea mays* ssp. *mays*) is adapted to a wide range of climatic and edaphic conditions. Here, we focus specifically on the potential role of root anatomical variation in this adaptation. In light of the investment required to characterize root anatomy, we present a machine learning approach using environmental descriptors to project trait variation from a relatively small training panel onto a larger panel of genotyped and georeferenced Mexican maize accessions. The resulting models defined potential biologically relevant clines across a complex environment and were used subsequently in genotype-environment association. We found evidence of systematic variation in maize root anatomy across Mexico, notably a prevalence of trait combinations favoring a reduction in axial conductance in cooler, drier highland areas. We discuss our results in the context of previously described water-banking strategies and present candidate genes that are associated with both root anatomical and environmental variation. Our strategy is a refinement of standard environmental genome wide association analysis that is applicable whenever a training set of georeferenced phenotypic data is available.

## INTRODUCTION

Abiotic stress is a major driver of plant phenotypic diversity (Lowry 2012), acting to select locally adapted varieties with specific morphological, physiological, and phenological traits (Fumagalli et al. 2011; Stebbins 1952; Hereford 2009https://paperpile.com/c/jHvuxa/s6Ua). Differences in such selective pressures over a continuously varying environment produce clines of genetic and phenotypic variation, reflecting the shifting costs and benefits of diverse biological strategies (Joswig et al. 2022). Although plants are plastic in the face of environmental challenges (Des Marais et al. 2013; Lasky et al. 2014), locally adapted specialists can constitutively express adaptive strategies, anticipating the need for the induction of stress responses (Levins 1968; von Heckel et al. 2016; Aguilar-Rangel et al. 2017). As a consequence, genotypes sourced from diverse locations will typically still display trait variation indicative of adaptation to their home environments when grown in a benign common garden (Stinchcombe et al. 2004; Janzen et al. 2022; Shimono et al. 2009).

Mexican native maize (*Zea mays* ssp. *mays*) varieties (“landraces”) represent an attractive system for the study of local adaptation. Mexico is the center of origin of maize, and today hosts 59 described native varieties cultivated from sea level to an elevation of 3,400 m, in environments ranging from semi-arid to hot and humid (Ruiz Corral et al. 2008; Arteaga et al. 2016; Perales and Golicher 2014). This diversity has been extensively sampled, and large collections of georeferenced and genetically characterized material are available (Arteaga et al. 2016; Romero Navarro et al. 2017; Mercer and Perales 2019; Janzen et al. 2022). Throughout maize domestication and diversification, farmers have consciously selected for agronomically and culturally desirable traits, principally targeting the female inflorescence (the ear) to produce a rich variety of form (Louette and Smale 2000; Bellon et al. 2018). In parallel, unconscious selection has likely acted to adapt varieties to local conditions (Romero Navarro et al. 2017; Mercer and Perales 2019; Janzen et al. 2022) and enhance tolerance to different environmental stressors after dispersal to new environments (Magalhaes et al. 2007;Eagles and Lothrop 1994; Bayuelo-Jiménez et al. 2011).

In this study, we focus specifically on the potential role of root trait variation in the adaptation of maize to different climatic and edaphic environments in Mexico. Although less easily visible than aboveground traits, the maize root system has been substantially impacted by domestication, diversification and modern breeding (Gaudin et al. 2014; Chen et al. 2022; Lopez-Valdivia et al. 2022; Burton et al. 2013; Ren et al. 2022). Roots are fundamental to plant water and nutrient acquisition, and play a key role in both wild and domesticated plants in determining performance under resource limitation (Wahl and Ryser 2000; Markesteijn and Poorter 2009; Ma et al. 2018). Within maize specifically, root trait variation among inbred breeding lines has been linked to performance differences under both water (Jaramillo et al. 2013; Bomfim et al. 2011; Schneider et al. 2020) and nutrient (Schneider, Postma, et al. 2017; Galindo-Castañeda et al. 2018) limitation. Extensive root trait variation has also been reported among native maize varieties (Burton et al. 2013), although the associated functional impact and possible adaptive roles remain to be fully characterized. Variation in the plant root system can be considered from the anatomy of individual roots to overall root system architecture (Jung and McCouch 2013; Lynch 2019), traits at all levels interact to determine overall root system function in the context of a given environment (Klein et al. 2020). Here, we limit ourselves to consideration of variation in root anatomy.

The maize root system consists of a variety of root classes that vary in function and importance during development (Atkinson et al. 2014; Viana et al. 2022; Hochholdinger 2009). Within this range, root anatomy develops on a basic pattern of radially organized tissue types: an external epidermis, the ground tissue, and an inner stele containing the pericycle and vasculature (Lynch et al. 2021). The epidermis protects the inner layers from physical damage and is in direct contact with the rhizosphere, playing a key role in water and nutrient exchange. The ground tissue is further differentiated into cortex and the endodermis that encloses the stele. In the model plant *Arabidopsis,* the cortex is composed of only two layers, a single layer of cortical parenchyma and the endodermis. In maize, however, the cortex divides to form multiple cell layers, impacting the physical (Chimungu et al. 2015), hydraulic (Heymans et al. 2020; Sidhu et al. 2023) and radial nutrient transport (Hu et al. 2014; Schneider et al. 2017) properties of the root, as well as accommodating beneficial endomycorrhizal fungi (Sawers et al. 2008; Bennett and Groten 2022). Developmental and environmental cues can trigger cells in the cortex to undergo programmed cell death and form aerenchyma. The resulting cortical air-filled lacunae help maintain gas exchange and mitigate hypoxia under flooding (Colmer 2003; Mano and Nakazono 2021). In addition, the reduction in root metabolic cost following aerenchyma formation can be beneficial in resource limiting conditions including drought and low availability of nitrogen or phosphorus (Jaramillo et al. 2013; Galindo-Castañeda et al. 2018). The cell walls of the endodermis are impregnated with suberin to form the Casparian strip and, in later development, further reinforced with lignin to act as a barrier that restricts apoplastic transport into the stele from the surrounding cortical root tissue. The central stele contains the xylem and phloem vessels that axially transport water and nutrients. The xylem is composed of small protoxylem vessels and larger metaxylem vessels, the latter providing the majority of the transport capacity in mature root tissues (Doussan et al. 1998). The size and number of metaxylem vessels and living cortical area influence root radial and axial hydraulic properties (conductivity and conductance; (Frensch and Steudle 1989; Schneider, Wojciechowski, et al. 2017), impacting water capture and plant performance (Richards and Passioura 1989; Couvreur et al. 2012).

In this study, we aim to characterize heritable variation in root anatomical traits in Mexican native maize and to associate patterns of phenotypic and genetic variation with the source environment. The genetic basis of local adaptation can be characterized through associations between genotype and phenotypes involved in local adaptation or between genotype and environment (Fournier-Level et al. 2011; Hoban et al. 2016). Approaches using genotype-environment association (GEA) have the advantage of not requiring resource-costly phenotypic characterization, although they must assume local adaptation to have occurred over the tested environments and can be complicated by the confounding effects of population structure (Lasky et al. 2023). One challenge of GEA is that it can be unclear which aspect(s) of the environment might drive selection, such that it may be unclear what associations to test. Here, we present an approach that first fits a trait relationship across the multivariate environment using a training set of phenotypically characterized, georeferenced varieties. This fitted model is then used to project trait values onto a larger set of available genotyped and georeferenced varieties, and the predicted trait values then used in genome-wide association (GWA) to find putative adaptive loci. We compare the results of our approach with standard GEA and a previously published phenotypic GWA using modern maize breeding lines. We present evidence for systematic variation in root anatomy driven by differences across the Mexican environmental landscape, identifying candidate genetic variants and linked genes associated with both phenotypic and genetic clines.

## RESULTS

### Root anatomy varies among Mexican native maize varieties

To characterize the relationship between root anatomy and environment in Mexican native maize, we assigned georeference data to 39 Mexican accessions phenotypically characterized in a previous root anatomy study (Burton et al., 2013; hereafter, the Burton panel) and extracted associated climate and soil descriptors from publicly available databases (see Materials and Methods). We supplemented published trait data with a re-analysis of the original cross-sectional images to generate a final phenotypic dataset of 16 anatomical traits (Table 1). We performed a principal component (PC) analysis on the phenotypic data: PC1 was negatively correlated with root cross-sectional area (Fig. S1; Fig. 1B). PC2 and PC3 captured allometric relationships among traits (Fig. 1A, B): PC2 was associated with variation in the relative contribution of the stele to the total area, with *total metaxylem vessel area* and associated metaxylem traits loading antagonistically to *total cortical area* and *root cross-section area*; PC3 captured an apparent trade-off between *cortical cell file number* and *cortical cell size* along with variation in cortical *aerenchyma area*. In addition to grouping traits by PC analysis, we used a modeling pipeline linking the GRANAR and MECHA packages to simulate cross-sectional anatomy (Heymans et al. 2020) and predict radial conductivity (*k_r_*) and radial and axial conductance (*K_r_*, *K_x_*; Couvreur et al. 2018; Fig. S2).

**Figure 1.**
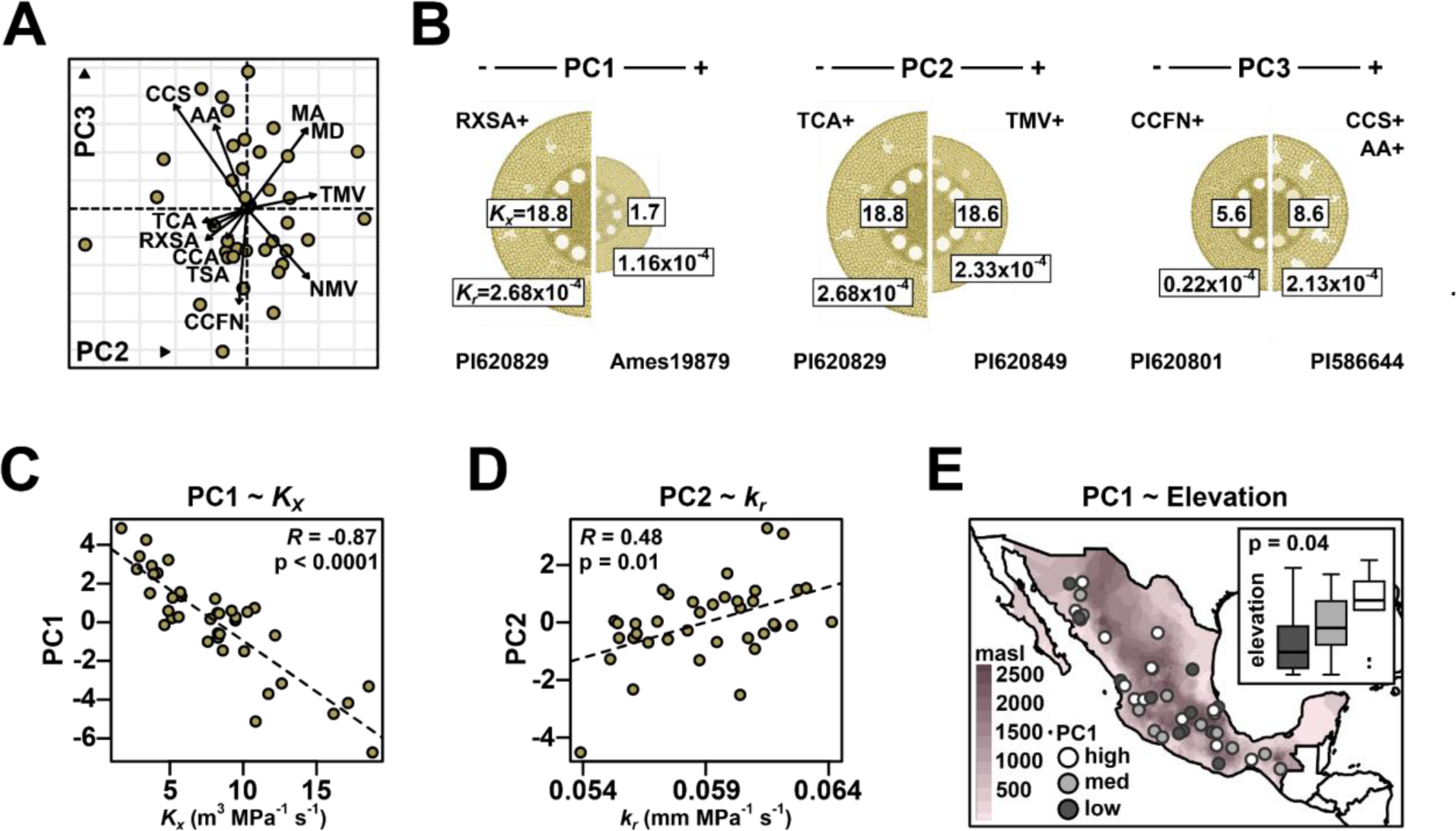
Root anatomy varies in Mexican native maize. A) PC2 and PC3 loadings for root anatomical traits of accessions from Burton et al. 2013. Trait description and codes as Table 1. B) Representative cross sections of extreme high- and low-loading individuals for PCs 1-3 rendered using GRANAR, scaled to the measured *root cross-section area*. Trait codes indicate broad trends seen in trait loading on the PCs. Boxed numbers adjacent to the central stele show modeled axial conductance (*K_x_*). Boxed numbers on the outer epidermis show modeled radial conductance (*K_r_*). Accession numbers are given at the base of the images. C) Correlation between modeled *K_x_* and anatomical PC1. D) Correlation between modeled radial conductivity (*k_r_*) and anatomical PC2. E) Accession source labeled by loadings on PC1, divided into terciles as low, medium (med) or high. Base map shaded by elevation. Inset box plots show the median and quartile elevation for the low, med and high PC loading groups. Whiskers extend to the most extreme points within 1.5x box length; outlying values beyond this range are shown as points. Stated p-value refers to an ANOVA for differences in elevation among the PC1 tercile groups.

**Table 1.**
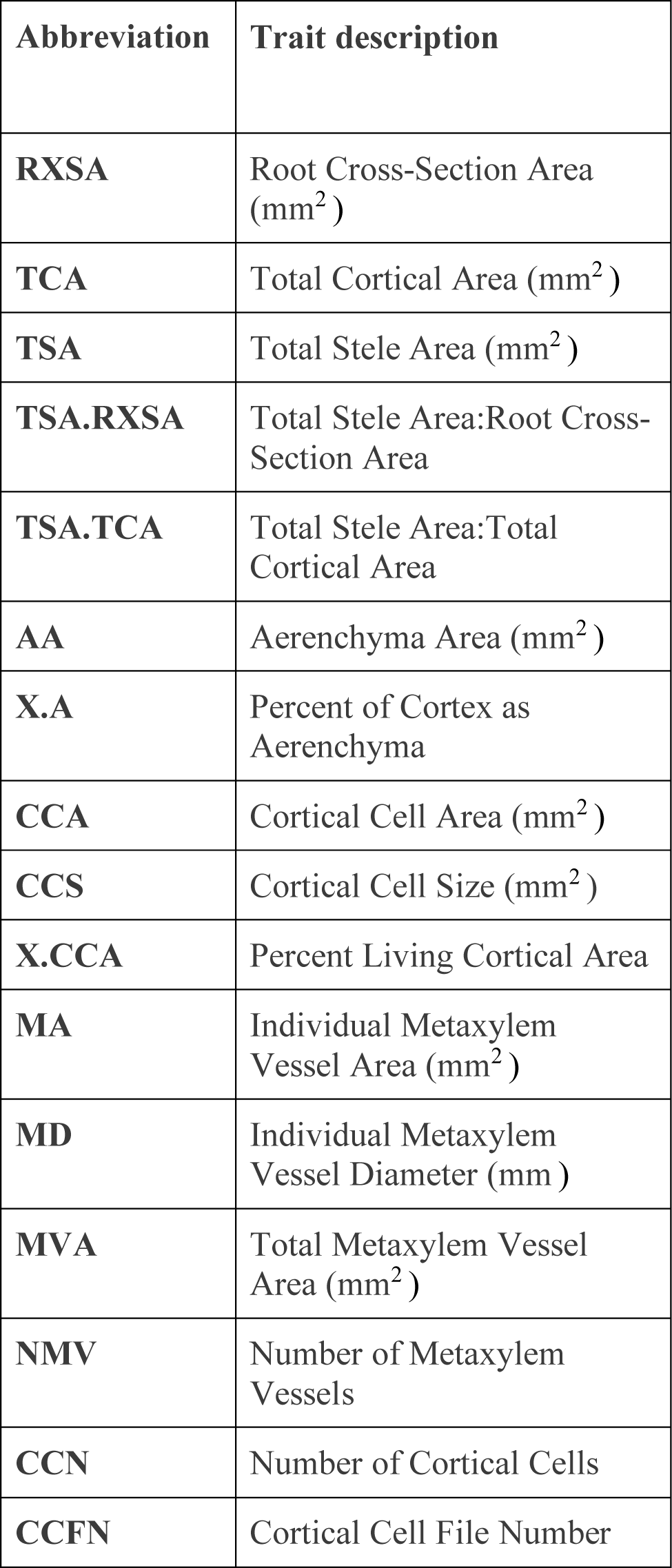
Description of root anatomical traits used in this study.

Axial and radial hydraulic conductance were negatively correlated with PC1, indicating the greater capacity of larger diameter roots for water transport (Fig. 1C). PC1 and PC2 were both positively correlated with radial conductivity (Fig. 1D), reflecting the greater ease of water transport across roots with less cortex (Heymans et al. 2020). Further associations between anatomical trait PCs and derived hydraulic properties were not easily captured by simple correlations (Fig. S2). As a first attempt to identify clinal relationships between root anatomy and local environment, we examined the correlation of root anatomy PCs to four basic environmental descriptors (elevation, annual precipitation, mean temperature, and soil pH). After adjusting p-values for multiple comparison testing, we did not find evidence for variation in root anatomy (Fig. S3; S4) or derived hydraulic properties (Fig. S5) to have significant associations to environmental descriptors of accessions’ point of origin. We did find mild evidence for root anatomical variation summarized by PC1 to be related to elevation (Fig. S3). When dividing PC1 axes into terciles, individuals with the most positive PC1 loadings were sourced from the highest elevations (Fig. 1E).

### Combined environmental descriptors predict variation in root anatomy

Local adaptation is driven by varied aspects of the environment and their interactions. To capture more complex trait-environment relationships, we used a feature-reduction method (Boruta algorithm; Kursa and Rudnicki 2010) to select the most informative of a full set of 157 available environmental descriptors for each anatomical trait, and subsequently combined the chosen descriptors into random forest (RF) models to relate environment and trait (Fig. 2A; S6). Nine of the 16 tested root anatomical traits were associated with environmental descriptors by the Boruta algorithm (Table S1). 39 different environmental descriptors were used as input for RF models across the 9 modeled anatomical traits, with individual models using from two (*percent of cortex as aerenchyma*) to 16 (*total metaxylem vessel area*) environmental descriptors. We observed varying goodness-of-fit from RF models and the R-squared for predicted vs observed trait values ranged from 0.32 (*aerenchyma area*) to 0.01 (*total metaxylem vessel area*) (Fig. S7).

**Figure 2.**
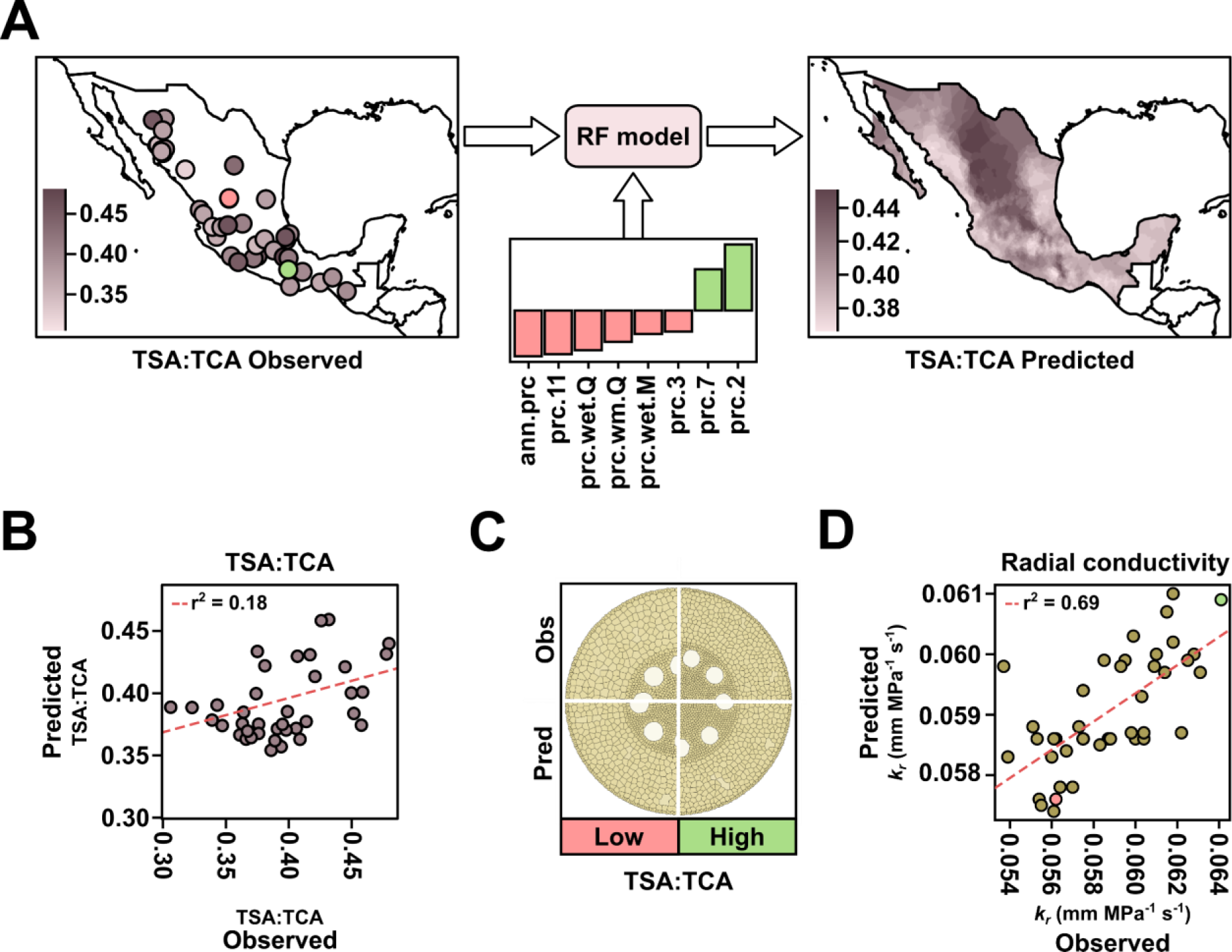
Home environment predicts root anatomy in Mexican native maize. A) Random forest (RF) modeling for the ratio of *total stele area:total cortical area* (TSA:TCA). Accession point of origin colored by observed TSA:TCA from Burton 2013. Individuals colored pink and green denote the accessions with the lowest (AMES19907) and highest (PI629263) observed TSA:TCA, respectively. Trait-specific significant environmental descriptors identified by the Boruta method used for RF model construction, displayed as SHapley Additive exPlanations (SHAP) contributions. Smoothed RF predicted TSA:TCA for native Mexican maize. B) RF predicted vs observed TSA:TCA values for all individuals used in model training and validation. C) Composite GRANAR representation of observed (obs) and predicted (pred) GRANAR sections for the accessions with the lowest (low, pink) and highest (high, green) observed TSA:TCA. Predicted GRANAR cross-sections use predictions for all traits for which RF models were constructed and are rendered at the same size. D) Predicted vs observed radial conductivity (*k_r_*) for all individuals used in model training and validation. Predicted *k_r_* was calculated using RF anatomical predictions and observed *k_r_* was calculated using observed anatomical values from Burton et al. 2013. The individuals with the lowest and highest observed TSA:TCA are colored pink and green, respectively. Dashed line is the coefficient of determination for all plotted points.

We used the GRANAR-MECHA pipeline to combine predicted trait values and compared observed and predicted anatomies, both graphically (Fig. 2C) and with respect to hydraulic properties (Fig. 2D). Modeled conductivity and conductance values for predicted anatomies correlated well with values from the observed data (radial conductivity, *r* = 0.69, p < 0.01; radial conductance *r* = 0.28, p = 0.07; axial conductance *r* = 0.59, p < 0.01; Fig. S8), indicating that our RF models successfully captured differences in anatomical traits that impact root hydraulic properties.

### Random forest prediction of root anatomy across Mexican native maize

To estimate root anatomical diversity across a broader sampling of native Mexican maize, we applied our Burton-trained RF models to a larger collection of 1791 genotyped and georeferenced Mexican accessions (hereafter, the CIMMyT panel; Romero Navarro et al. 2017; Fig. S9). We used the georeference data to link environmental descriptors to each accession and passed these to the RF models, generating a complete phenotypic set of 9 estimated root anatomical traits for the 1791 accessions (Supplementary Information). To summarize patterns among the predicted trait values, we used partition-against-medians (PAM) clustering (Klein et al. 2020; Maechler et al. 2021) to group the accessions into seven phenotypic clusters (Fig. S10, S11). The clusters 1 through 7 were composed of 308, 366, 370, 231, 158, 277 and 131 accessions, respectively. The structure defined by the clustering was not strong (mean silhouette value = 0.25), reflecting the continuous nature of the environmental descriptors driving the RF models, but did provide a context for subsequent analyses. We also passed the median trait values of each cluster to the GRANAR-MECHA pipeline to obtain average anatomies and hydraulic properties.

Clusters were distinguished by the relative elaboration of cortex and stele and associated hydraulic properties (Fig. 3). In Clusters 1 and 3, the stele (*total stele area: root cross-section area*; *total stele area: total cortical area*) was relatively small, although individual metaxylem vessels were large (*individual metaxylem area*, *individual metaxylem diameter*) and, consequently, the *total metaxylem vessel area* and axial conductance were relatively high. In contrast, Clusters 5, 2, and 6 were distinguished by a small stele and small metaxylem vessels, associated with low axial conductance relative to other clusters. In Cluster 5, the small size of the metaxylem vessels was further associated with a low *number of metaxylem vessels* resulting in the lowest *total metaxylem vessel area* and the lowest axial conductance of the clusters.

**Figure 3.**
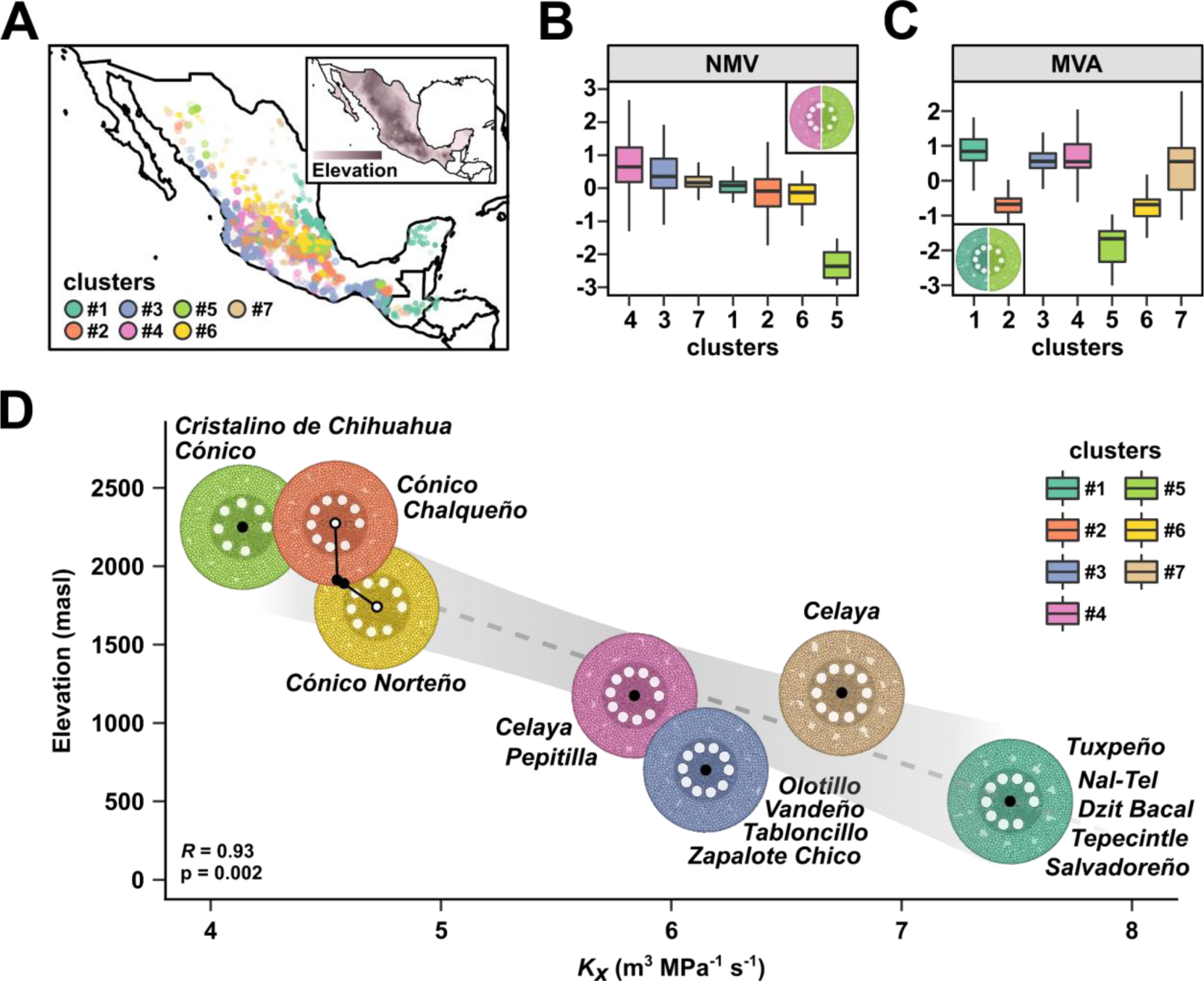
Groups defined by shared root anatomical characteristics originate from distinct environments. 1791 maize accessions forming the CIMMyT panel were grouped into seven clusters based on eight RF predicted root anatomical traits. A) Geographical distribution of the clusters. Inset shows elevation, with darker shading corresponding to higher values. B) Centered and scaled *Number of metaxylem vessels* (NMV) and C) *Total metaxylem vessel area* (MVA) in the seven clusters. Inset shows composite GRANAR representation generated from the median trait values of highest (left) and lowest (right) scoring clusters. D) Variation in mean cluster predicted axial conductance across elevation. Black points indicate the mean axial conductance calculated using RF predicted anatomy vs the mean elevation at the point of collection for each of the seven clusters. For clusters 2 and 6, root-cross section renderings are plotted off the regression line for easier view of digitized anatomy. Root cross-section images are GRANAR representations generated using the median trait values for each cluster, all rendered at the same size. Native varieties overrepresented in each cluster are listed adjacent to the GRANAR images.

We examined the clusters with respect to previous morphological-isozymatic (Sanchez G. et al. 2000) and environmental (Ruiz Corral et al. 2008) classifications of Mexican native maize (Table S2, S3; Fig. S11). Following trends reported for overall Mexican maize diversity (Sanchez G. et al. 2000), our clusters were structured with respect to elevation (Fig. 3). Clusters 2, 5 and 6 were enriched (Fisher test for variety count in or out of the cluster, adj.p < 0.05) for varieties belonging to the previously defined highland group (Sanchez G. et al. 2000; Fig. 3; Table S3). Cluster 5, containing the highest elevation varieties, was centered on Mexico City, although it also contained accessions from the highlands of Chihuahua in northern Mexico; Cluster 6 extended from north to south along the Sierra Madre Occidental; Cluster 2 was again centered on Mexico City, although with greater representation further west along the trans Mexican volcanic belt than Cluster 6 and included several accessions from the Chiapas highlands on the southern border of Mexico. The mid-elevation Clusters 4 and 7 were loosely sourced from the center-to-south and center-to-north of Mexico, respectively. The Clusters 1 and 3 were enriched for varieties in the lowland short-to-medium maturity and tropical dent groups (Sanchez G. et al. 2000; Table S2, S3), with Cluster 1 from the Gulf Coast, the Yucatan and lowland Guatemala and Cluster 3 from the Pacific Coast. Prior environmental classification was in line with the observed elevational cline: Clusters 2, 5 and 6 were enriched for varieties previously assigned to “temperate to semi-hot” environments; Clusters 1 and 3 were enriched for varieties assigned to the “very hot” niche (Ruiz Corral et al. 2008; Fig. S12; Table S3).

Considering the average anatomies and mean values of environmental descriptors associated with each cluster, we could discern a broad trend of a reduction in axial conductance with increasing elevation (Fig. 3D). Our models associated the colder, drier highland niche (>2,500 masl; Eagles and Lothrop 1994; Ruiz Corral et al. 2008) with both fewer and smaller metaxylem vessel elements (Cluster 5). Conversely, the hot, wet lowlands (Eagles and Lothrop 1994; Ruiz Corral et al. 2008) were associated with a relatively larger stele accommodating a greater number of larger metaxylem vessel elements (Clusters 1 and 3).

### Novel phenotypic evaluation supports random forest predicted variation in axial conductance

To empirically evaluate our RF models, we characterized eight maize accessions from across Mexico that had not been used previously in the Burton study (Fig. 4A, B). We measured root anatomical traits following the Burton protocol, and for each trait compared the observed best linear unbiased predictor (BLUP) with the results of our environmental RF predictions (Fig. S13). The correlation between observed BLUPs and RF predictions ranged from relatively high for cortical traits (*aerenchyma area*, *r* = 0.71; *percent of cortex as aerenchyma*, *r* = 0.65; *percent of cortex as cortical cells*, *r* = 0.52) to lower for metaxylem vessel traits (*number of metaxylem vessels*, *r* = 0.30; *total metaxylem vessel area, r* = 0.27; *individual metaxylem vessel area*, *r* = 0.11) and allometric traits (*total stele area:root cross-section area, r* = 0.18*; total stele area:total cortical area r* = 0.06). Predictions of *individual metaxylem vessel diameter* were not well supported by observed values (*r* = -0.24).

**Figure 4.**
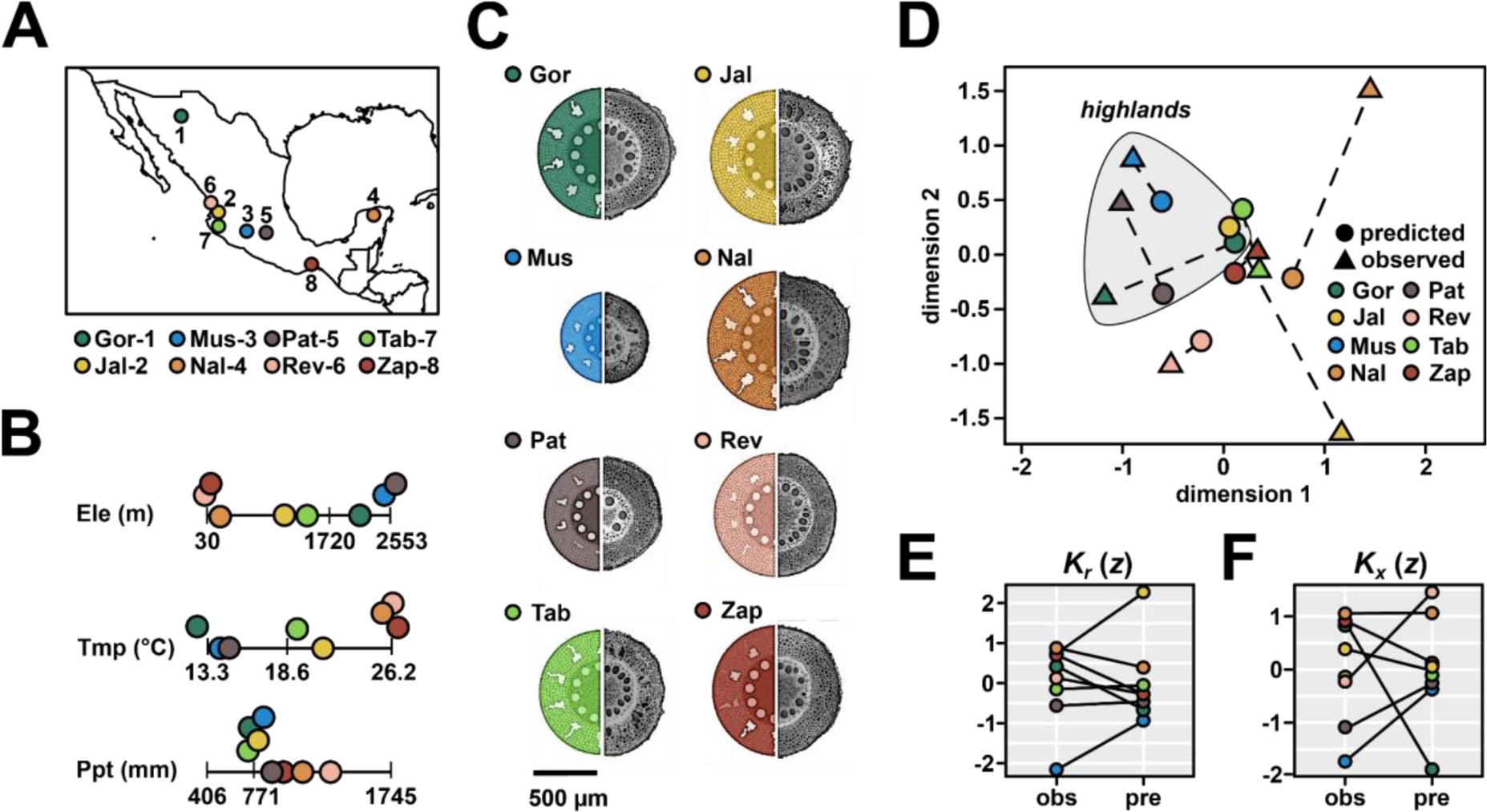
Novel phenotypic data is consistent with model predictions. A) Source of eight selected native maize varieties: 1) Gordo, 2) Jala, 3) Mushito, 4) Nal Tel, 5) Palomero Toluqueño, 6) Reventador, 7) Tabloncillo, 8) Zapalote Chico. B) Elevation (Ele), annual mean temperature (Tmp), and annual precipitation (Ppt) at source locality for the eight native varieties. Bars on the line plots represent the 5%, 50%, and 95% quantiles for each environmental descriptor across all CIMMyT panel individuals included in this study (1791). Points color-coded as A. C) GRANAR renderings of observed anatomical BLUPs across node two and three roots (colored) and photographs of representative cross-sections of third-node roots, scaled to the mean measured *root cross-section area.* D) Procrustes analysis comparing distribution of RF-predicted (circles) and observed (triangles) anatomical traits. For each variety, predicted and observed projections are linked with a dotted line. An arbitrary ellipse was added around three varieties sourced from high elevation. E) Comparison of standardized observed (obs) and predicted (pre) values of modeled radial conductance (*K_r_*). Values for each variety are connected to illustrate the level of consistency in ranking. F) as E, showing modeled axial conductance (*K_x_*).

We assessed overall concordance between observed and predicted anatomy by using the Procrustes transformation (Schönemann 1966) to minimize the distance between each set of observed and predicted trait values across the eight accessions. Observations and predictions were well matched for six of the eight accessions, with the Jala and Nal Tel accessions being a poorer fit (Fig. 4D; Fig. S14). The difference in overall root anatomy in material sourced from the highlands and lowlands was well supported by both observed and predicted trait values (Fig. 4D, S15) and modeled hydraulic properties (Fig. 4E, F). Overall, although individual traits were not always well predicted for any given accession, our methodology and training data were sufficient to capture broad stratification of anatomical traits and hydraulic properties across the environment.

### Genome wide association analysis using predicted trait values identifies novel candidate genes

To look for genetic evidence linking root anatomy to the local environment in Mexican maize, we ran a GWA analysis on the CIMMyT panel using the RF predicted trait values. Given the nature of the RF models, this prediction GWA is, in effect, a development of a standard environmental GWA analysis with modeled trait values capturing complex combinations of the individual environmental descriptors used in RF model construction. For comparison, we ran separate environmental GWA analyses for each of the 39 environmental descriptors used in RF modeling, and also re-analyzed published phenotypic data for a panel of 175 maize inbred lines (hereafter, the WIDP panel, Schneider et al. 2020). We extracted phenotypic data for 8 of our 9 RF modeled traits (not including *percent of living cortical area*), combining values obtained for well-watered and water-limited treatments into a single GWA model (Runcie and Crawford 2019), estimating variant main (G) and variant x treatment (GxE) effects. To facilitate comparison across panels genotyped using different platforms, we used the MAGMA pipeline (de Leeuw et al. 2015) to combine signals across single nucleotide polymorphisms (SNPs) to a single gene level value. Here, we assigned any SNP +/- 2.5 kb from an annotated gene model to that gene. In the following discussion of overlap between our different GWA analyses, we consider only genes captured in both CIMMyT and WIDP markersets. In later identification of the genes of greatest interest from the predicted GWA analysis, we do not take the WIDP markeset into account.

To compare CIMMyT “predicted”, CIMMyT “environment”, WIDP “G” and WIDP “GxE” GWA analyses, we selected the top 100 genes (determined by p-value) per trait for each analysis and combined these into candidate gene lists, obtaining sets of 636 unique “prediction” genes, 1,282 unique “environment” genes, 542 unique “G” genes and 576 unique “GxE” genes (Fig. 5; Supplementary Information). Only 19% of the prediction genes were also present in the environment set (Fig. 5B), indicating that the two were not redundant and that the prediction set was capturing patterns not revealed by separate analyses of the individual environmental descriptors. For example, a region of the short arm of chromosome 10 was linked to *mean metaxylem vessel diameter* in the prediction analysis (Fig. S16). In this case, the -log_10_(P) value of the most significant SNP for the predicted trait is approximately double that of the best supported environmental descriptor (precipitation in October). The best supported SNP in this region fell within the gene *Trichome birefringence-like 10* (*Tbl10*; Zm00001d023378; Fig. S16). Natural variation in *Tbl10* has previously been linked to variation in flowering time (Chen et al. 2012; Kusmec et al. 2017), height (Wang et al. 2022), and root diameter (Pace et al. 2015). As such, *Tbl10* illustrates a compelling candidate for further follow-up that would not have been identified by standard environmental GWA.

**Figure 5.**
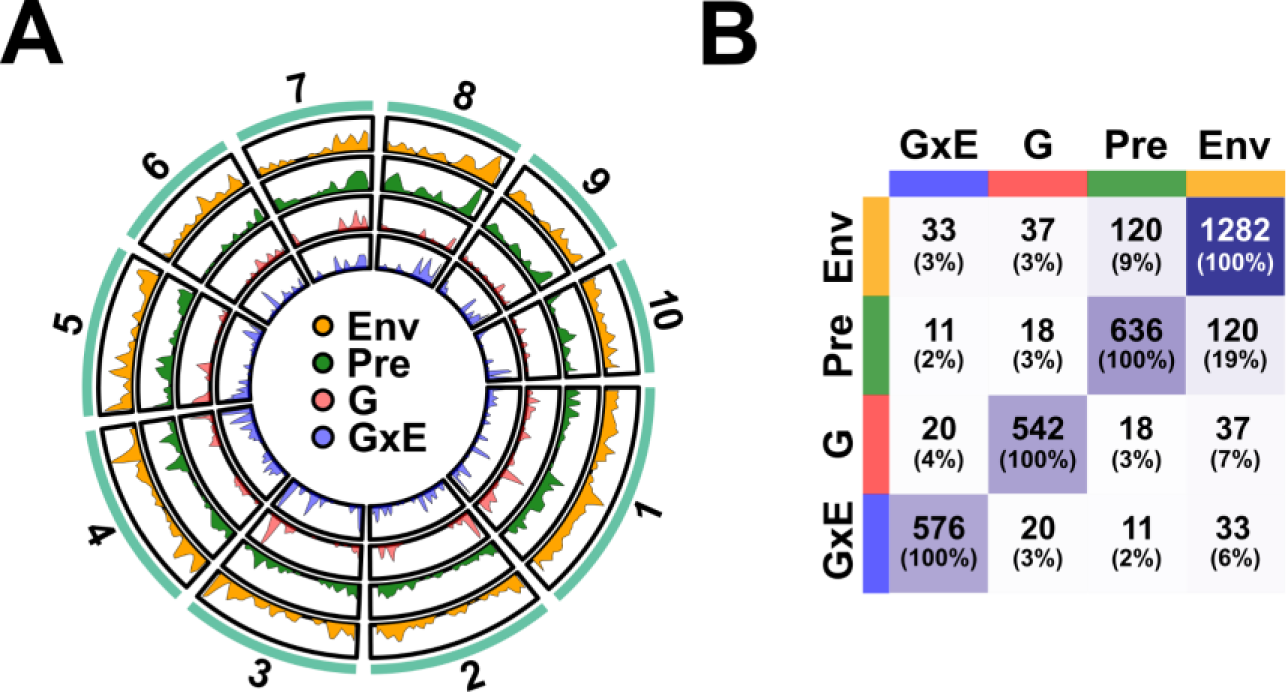
Evidence for a shared genetic basis of root anatomical variation between inbred breeding lines and Mexican native maize. A) Distribution of top 100 genes from GWA analyses of WIDP and CIMMyT accessions across the genome. The shaded area represents the density of top genes overlapped with window regions (1 x 10^7^ bp). Sector names represent the number of chromosomes. B) The number of pairwise overlapping genes among the GWA gene sets. The darker the color of the squares indicates a higher number of genes. Totals in parentheses show the percentage of the row set in the other sets.

### The gene *Vq29* is linked to variation in both metaxylem traits and source elevation

There was no evidence that the prediction set was enriched for WIDP root anatomy candidate genes with respect to the environment set - both contained 2-3% WIDP G and GxE genes (Fig. 5B). Nonetheless, we do consider the 29 genes identified in both prediction and WIDP (G and/or GxE) GWA to be high confidence candidates for further characterization (Fig. 5B, S17; Table S4). For example, the gene *Vq29* (Zm00001d015397) on the short arm of chromosome 5 was associated with *number of metaxylem vessels* and *total metaxylem vessel area* in the prediction GWA and with *individual metaxylem vessel area* and *individual metaxylem vessel diameter* in the WIDP G analysis (Fig. 6A). The *Vq29* gene is predicted to encode a VQ domain transcription factor, part of a large family of proteins that interact with members of the WRKY family under stress (Song et al. 2015), including in response to hypoxia, ozone or nitric oxide (León et al. 2020). The minor allele of the highest scoring SNP (S5:88306863) was associated with greater *number of metaxylem vessels* and *total metaxylem vessel area* and declined in frequency within our clusters with increasing predicted values of these same traits (Fig. 6B). Based on gene expression atlas data (Walley et al. 2016), *Vq29* is most highly expressed in the roots, consistent with a role in metaxylem development (Fig. 6C). Geographically, the minor allele of S5:88306863 was most prevalent in the central Mexican highlands (Fig. 6D), and the MAF increased with mean elevation across our previously defined clusters (Fig. 6E). In summary, *Vq29* nicely illustrates an example of a candidate gene associated with phenotypic variation in root anatomy in the inbred WIDP panel that also shows clinal genetic variation across the Mexican environment.

**Figure 6.**
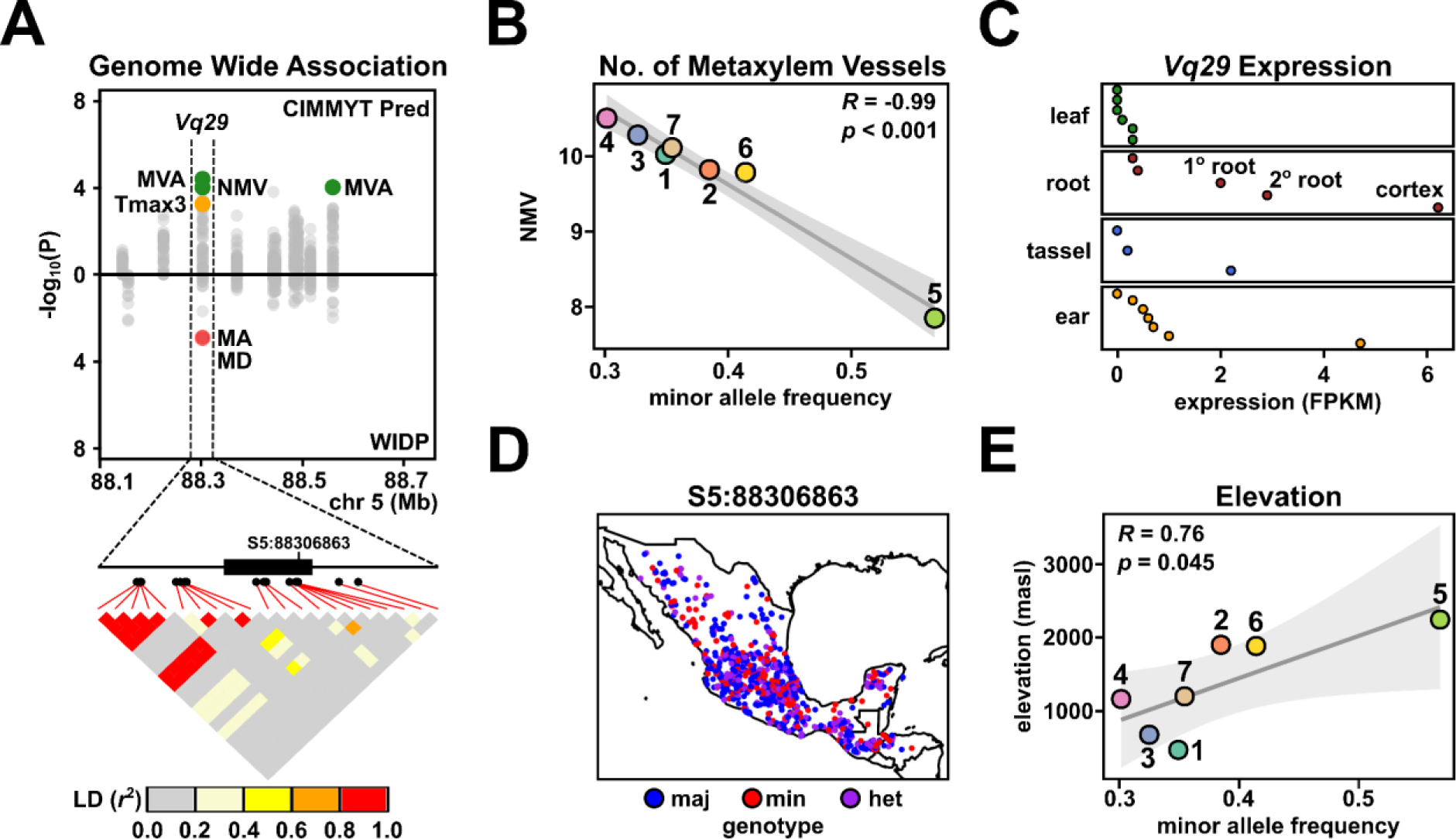
The gene *Vq29* is linked to variation in root anatomy and source environment. A) Miami plot showing GWA support (-log_10_P) for association with root anatomy for genes on a region of chromosome 5. Points above the x-axis show support for predicted phenotypes in the CIMMyT panel; points below the x-axis show support for observed phenotypes for the WIDP panel. The gene *Vq29* is associated with *total metaxylem vessel area* (MVA), *number of metaxylem vessels* (NMV), *individual metaxylem vessel area* (MA) and *individual metaxylem vessel diameter* (MD) across the two analyses. Image below the Miami plot shows the *Vq29* gene model (CDS as filled box), SNP position (filled circles) and pairwise linkage disequilibrium (LD). The position of the focal SNP S5:88306863 is highlighted. B) Correlation between frequency of the minor allele at S5:88306863 in the previously defined CIMMyT clusters and mean predicted NMV. C) Expression of *Vq29* in four named tissues from publicly available expression data. Points show different subsamples. The root cortex, corresponding to the highest expression, is highlighted. D) Geographic allele-distribution of S5:88306863 in the CIMMyT panel. E) Correlation between frequency of the minor allele at S5:88306863 and mean elevation at source in the CIMMyT clusters.

## DISCUSSION

We have presented evidence that variation in root anatomy contributes to local adaptation in Mexican native maize. We used predictive models to define biologically relevant clines over which we identified both genotypic and phenotypic variation. Shared GWA candidates between Mexican native maize and modern inbred lines indicated an element of common genetic architecture, although we also identified novel candidates specific to the native Mexican material. Phenotypic patterns suggested that local differences in precipitation and temperature are associated with heritable variation in maize root anatomy. Root anatomical variation broadly followed the established grouping of Mexican maize varieties, themselves strongly stratified by environment. Our observations are consistent with a role for root anatomical variation in local adaptation. Our combination of environment-based models and GWA allowed us to leverage a relatively small sample of phenotypically characterized locally adapted varieties to identify novel associations between phenotype, genotype and environment in the context of broader maize diversity.

Our analyses highlight a predominance of anatomies predicted to reduce axial conductance in material sourced from arid subtropical or temperate environments. In both observed and predicted phenotypic data, varieties from the cooler, drier highland regions were associated with fewer and/or narrower metaxylem vessels and a reduction in the area of the stele with respect to cortex. Comprehensive revision of data across taxa has previously suggested that the capacity for axial water transport is typically greater in plants from wet environments and reduced in plants adapted to xeric conditions (Feng et al. 2016; Lynch et al. 2021). Although somewhat counterintuitive, reducing water uptake under dry conditions may benefit plants by reducing root tip desiccation (Richards and Passioura 1989), preventing cavitation (Nardini et al. 2013) and enabling the conservation of soil water resources across the growing season (Richards and Passioura 1989; Leitner et al. 2014). Studies of interspecific variation in crops support these hypotheses with both narrower metaxylem vessels (Priatama et al. 2022; Allah et al. 2010; Purushothaman et al. 2013; Peña-Valdivia et al. 2005) and fewer metaxylem vessels (Strock et al. 2021) being associated with enhanced drought tolerance. Similarly, selection for reduced xylem vessel diameter in Australian wheat has been reported to successfully increase yield under water limitation (Richards and Passioura 1989). In the Mexican highlands, farmers traditionally plant prior to the beginning of the annual rains to maximize the length of the growing season and ensure crops reach maturity prior to the first frosts (Eagles and Lothrop 1994). As a consequence, seed is deep planted to better access residual soil moisture, as well as to offer protection from low temperatures, a practice also employed in the southwestern US (Collins 1914). For wheat varieties reliant on residual soil moisture during early growth, reduced root conductance has been correlated with increased yield (Passioura 1972). Mexican highland maize may similarly benefit from rationing water use early in the season (Fischer et al. 1983; Hayano-Kanashiro et al. 2009) with reduced axial conductance contributing to this adaptive water saving strategy.

We observed variation in the formation of root cortical aerenchyma, with varieties sourced from regions of higher precipitation generally being associated with greater aerenchyma formation. Root cortical aerenchyma forms constitutively in wetland crops such as rice and in maize wild relatives endemic to regions of high precipitation (Mano et al. 2007; Mano and Nakazono 2021). Many cultivated maize genotypes lack constitutive aerenchyma; however, aerenchyma formation can be induced by environmental stresses, such as hypoxia (Yamauchi et al. 2016), drought (Zhu et al. 2010), heat (Hu et al. 2014) or nutrient starvation (Saengwilai et al. 2014; Galindo-Castañeda et al. 2018). Although greenhouse evaluation was conducted in benign condition, substantial aerenchyma production was observed (11% Jala, this study; 16% PI586644 in Burton et al. 2013) of the total cortical area in individual sections. In the field, aerenchyma plays a role in oxygenation of the root tissue under hypoxia (Jackson et al. 1985; Colmer 2003), while, in resource-limited conditions, the reduction in root metabolic cost resulting from aerenchyma formation may enhance the efficiency of foraging in terms of carbon invested (Klein et al. 2020; Lynch et al. 2021). On the other hand, with fewer living cortical cells a plant may be less able to accommodate mutualistic arbuscular mycorrhizal fungi, although the relationships between root anatomy, microbial interactions, environment and cortical burden remain to be fully understood (Saengwilai et al. 2014; Galindo-Castañeda et al. 2018; Strock et al. 2019). In our predictive analysis high *aerenchyma area* was associated with the varieties from the Gulf coast, the Yucatan (exemplified by Nal Tel in our greenhouse evaluation) and the region around the southern Mexican border extending into Guatemala. This last region covers the native range of the flooding tolerant teosinte *Zea mays* ssp. *huehuetenangensis* (Mano et al. 2005) and suggests flooding may have also exerted a selective pressure on the endemic maize.

The functional impact of root anatomical variation is contingent on root system architecture and, indeed, overall plant phenology (Lynch 2019). Differences in growth angle and branching determine the deployment of roots across the soil profile and will interact, synergistically or antagonistically, with root anatomy to impact overall root function. For example, the water-banking effect of reduced axial conductance discussed above has been shown to be enhanced in the context of a shallow root system architecture, enhancing the performance of inbred maize under drought (Strock et al. 2021). While there is a scarcity of information concerning root system architecture in Mexican maize, the limited data reveal remarkable structural diversity, indicating strong spatio-temporal variation in soil exploration (Heymans 2022). It has been noted that native maize root systems tend to be generally shallower compared to those of inbred lines (Burton et al. 2013; Ren et al. 2022). Interestingly, the highland varieties we found associated with reduced axial conductance have previously been described to have a high tendency to lodge (fall over) due to “poorly developed” root systems (Wellhausen et al. 1952). In practice, traditional management involves pilling of earth around the growing plant, freeing the root system from the need to provide mechanical support and perhaps allowing an overall reduction in root system development that contributes to water-banking.

In summary, our analyses indicate that reported variation in Mexican native maize root anatomy is distributed systematically over the environment, consistent with a role in local adaptation. We propose that predictive models based on a set of “signpost” accessions can define biologically relevant clines though complex environments, providing the appropriate axes against which to identify both phenotypic and genetic trends. Significantly, we obtained candidate genes from our predicted trait GWA that were not identified in one-by-one analyses of environmental descriptors, including candidates whose functional role in root anatomy is supported by previous studies of inbred maize (Fig. 5; Klein et al. 2020; Schneider et al. 2020). The combined use of field evaluation and *in silico* modeling has allowed great progress to be made in defining the functional impact of root anatomical variation (Heymans et al. 2020; Lynch et al. 2021; Sidhu et al. 2023). The further study of locally adapted native varieties has the potential to complement these other approaches. The history of native crop diversity is a natural experiment that has run for thousands of years, selection imposed by environmental conditions being integrated over many generations. As such, subtle signals that can be hard to detect in experimental evaluation may be amplified and detected as patterns of GEA.

## MATERIALS AND METHODS

### Phenotyped Burton panel

Phenotypic data from previous characterizations of greenhouse-grown native Mexican maize were obtained from Burton et al. 2013. After filtering for accessions of Mexican origin, subsequent analyses were completed with data from 39 georeferenced individuals. Additional root anatomical features including *total metaxylem vessel area* (MVA), *individual metaxylem diameter* (MD), *individual metaxylem area* (MA), *number metaxylem vessels* (NMV), and *cortical cell size* (CCS) were measured from the original Burton et al. cross-section images generated using *RootScan* v2.4, an imaging software designed to measure anatomical features of root cross-sections from digital images (Burton et al. 2012).

### Genotyped CIMMyT FOAM panel

Genotypes from a collection of 1791 native Mexican maize accessions from the CIMMyT Maize Germplasm Bank (CIMMyT panel) were obtained from (Navarro *et al*. 2017; Gates *et al*. 2019). In brief, sequences were generated using an Illumina HiSeq, and genotypes were called in TASSEL. Missing SNPs were imputed using BEAGLE4, and SNPs were further filtered for minor allele frequency >1%. The genotype data was uplifted to coordinates on the B73 v4 reference genome using Crossmap.

### Environmental Data

We compiled climatic and soil data for each representative of the long-term averages experienced by an accession’s point of origin for both the Burton and CIMMyT panel. All data used was sourced from publicly available sources with global coverage. Climate data was extracted using R/raster::extract (Hijmans 2023) following the methods described in (Lasky et al. 2015). Briefly, the first set of climate variables come from WorldClim and include information on monthly minimum, maximum, and mean temperatures; mean monthly precipitation; and other derived parameters of biological importance that take into account temperature and precipitation dynamics (Hijmans et al. 2005). Monthly and annual average potential evapotranspiration (PET), and a measure of aridity (mean annual precipitation divided by mean annual PET) that is calculated from WorldClim data were collected from the CGIAR-CSI Globality-Arbitdty database (Zomer et al. 2008). Information on inter-annual variability in precipitation, which may representative of areas where drought acclimation is important (Lasky et al. 2012) were calculated with data from the NCEP/NCAR Reanalysis project (https://psl.noaa.gov/data/reanalysis/reanalysis.shtml; Kalnay et al. 1996). Inter-annual variability in precipitation was obtained by calculating each calendar month’s coefficient of variation (CV) across years for each month’s surface precipitation rate. Information on estimated photosynthetically active radiation (PAR) for each quarter were averaged for data collected from NASA SRB (https://asdc.larc.nasa.gov/project/SRB).

Dynamics of evaporative demand on individuals in the form of vapor pressure deficit (VPD), or the difference between partial pressure of water vapor and maximum potential pressure was collected from the Climate Research Unit (New et al. 2002). In addition to climate data, we also included edaphic chemical and physical properties representative of the long term averages experienced by accessions. The soil data was collected from two sources, SoilGrids (Hengl et al. 2017) and the Global Soil Dataset (GSD; Shangguan et al. 2014). Data from GSD includes soil features of the topsoil and 1 meter below the surface. We found high concordance of values for topsoil and 1 meter below the surface and excluded the topsoil data from our dataset. All soil variables were cleaned by removing outliers and imputed missing values using the MICE package (van Buuren and Groothuis-Oudshoorn 2011; Fox et al. 2017).

### GRANAR representations and MECHA estimation of emergent hydraulic properties

Generator of Root Anatomy in R (GRANAR; https://granar.github.io), and the model of explicit cross-section hydraulic architecture (MECHA; https://mecharoot.github.io) are open-sourced computational tools. The first tool uses anatomical parameters as inputs to generate digital root anatomies. Once constructed, GRANAR root anatomies can be used for digital visualizations of anatomical parameters, and the anatomical network can be written as a XML file with the same format as CellSet output (Pound et al. 2012). The second tool uses the anatomical networks, such as the one generated by GRANAR to estimate emergent hydraulic properties. We used GRANAR to reconstruct virtual anatomies for all observed accessions of the Burton panel, our predictions of the Burton panel Mexican lines, predicted CIMMyT clusters, and predicted and observed novel germplasm grown in this study. For all aforementioned individuals, we estimated the root hydraulic conductance (*K_r_* and *K_x_*) and conductivity (*k_r_*) with MECHA. The subcellular hydraulic parameters are the same as in (Heymans et al. 2020), and the chosen hydraulic scenario accounts for the hydrophobic structures of an endodermal Casparian strip. The script used is available on a GitHub repository HydraulicViper/RootDiversity (doi: 10.5281/zenodo.10104521) under a GPL-3 license.

As not all anatomical features required for the GRANAR-MECHA pipeline were predictable with our RF models, a few transformations were required for predicted anatomical data to be input into GRANAR. With the exception of traits where RF predictions could directly be used as inputs (*number of metaxylem vessels, metaxylem vessel area*, *aerenchyma area*), we used constant values of the mean Burton panel (*root cross-section area*) or extrapolated values from RF predictions (*total stele area* calculated from RF predicted *total stele area:root cross-section area* using the Burton panel *root cross-section area* mean).

### Principal component analysis/initial trait/env association

We used principal component analysis (PCA) to initially explore internal anatomical variation captured in the greenhouse grown Burton panel. Phenotypes were constrained to only include “pure” traits, excluding proportional and percentage traits which are likely redundant and may be representative of more emergent properties. Principal components were built with R/ade4::dudi.pca and visualized with R/factoextra. We first compared phenotypic PC axes to calculated hydraulic properties to explore the combinations of anatomical traits that are related to variation in hydraulic properties. As an initial attempt to identify relationships between root anatomical traits and environmental variation, we compared the first three phenotypic PCs loadings and calculated hydraulic properties to core environmental features of the accessions’ point of origin (elevation, annual precipitation, annual mean temperature, soil pH). For all environmental, hydraulic, and phenotypic PC correlations, p-values were adjusted for multiple testing using the Holm method with R/stats::p.adjust (α = 0.05). As correlation between both root anatomical traits and derived hydraulic properties and environment of origin were weak, we then considered if combined environmental features were more able to describe variation in root anatomical traits and hydraulic properties.

### Feature Selection

We sought to determine if aspects of accessions’ home environment predict variation in root anatomical traits using a machine learning approach. Importantly, not all variables in the environmental dataset are related to root anatomical variation and not all tested root anatomy traits are significantly associated with variation in environmental features. Feature selection was employed to identify the anatomical traits that had relationships with environmental features (“environmentally related traits” from here on) and the environmental descriptors that described variation in those traits through eliminating unimportant variables. We used the function R/Boruta::boruta to obtain all important and tentatively important environmental features for each trait in our dataset (Kursa and Rudnicki 2010). Root anatomical traits which had significant variation described by at least two environmental features were considered environmentally related and retained for further analysis.

### Random Forest Models

We employed random forest (RF) to determine if variation in response variables (observed anatomical traits of the Burton panel) could be described by several explanatory variables (feature-selected environmental descriptors). For each environmentally related trait, we built a RF model that summarized how trait values are predicted to change across feature-selected environmental space. RF is a non-parametric classification method that constructs decision trees using subsets of input data to select the predictor variables that limit variance for the response variable predictive model (Breiman 2001). RF models have been found to have high predictive performance when tested data include a large number of predictor variables with little to no relationship to response variables (Fox et al. 2017); however, best practice is to limit the number of predictor variables to avoid model overfitting. As such, we limited response variables for each environmentally related trait model to be the specific environmental features identified in the Boruta feature selection step. As the environmental variables used in this study were continuous, RF models were built as regression trees. RF models were built using R/randomForest::randomForest, 5000 trees were built per model and one third the number of explanatory variables were tried at each split (Liaw and Wiener 2002). We increased our number of trees from the default value (500), to account for models with a large number of predictors and for increased stability of variable importance. Model success was evaluated with the percent variance explained output extracted from the randomForest package and the correlation coefficient between observed and RF predicted trait values. To determine the contribution of each boruta-identified environmental descriptor for constructed RF models, we calculated SHapley Additive exPlanations (SHAP) values (Lundberg and Lee 2017).

### Predicting traits using environmental relationships

Using the constructed RF models, unknown phenotypes of 1791 georeferenced, genotyped CIMMyT accessions (CIMMyT panel) were predicted from environmental descriptors of accession point of origin. Using R/caret::predict (Kuhn 2021), values for all nine environmentally associated traits were predicted for the 1791 CIMMyT panel. For each trained RF model, the full environmental dataset summarizing the source environment of the CIMMyT panel was constrained to include the environmental descriptors used to train each RF model. These constrained environmental descriptors were used as inputs in each trained RF model to calculate environmentally related traits for all individuals of the genotyped panel.

### Clustering analysis

CIMMyT FOAM accessions were clustered by predicted root anatomical traits using Partitioning Around Medoids (PAM) as previously described (Klein et al. 2020). Briefly, anatomical trait values were centered and scaled using R/caret (Kuhn 2021) and outlying values (> 3 standard deviations from the mean) removed. Within cluster sums of squares (WSS) were visualized as a function of cluster number using R/factoextra::fviz_nbclust (Kassambara and Mundt 2020). From inspection of the resulting curve, the accessions were grouped into seven clusters using R/cluster::pam (Maechler et al. 2021) under default settings. Primary variety (landrace) designations were assigned using data available from CIMMyT (www.mgb.cimmyt.org; 1454 accessions assigned), and these matched to existing morphological-isozymatic (Sanchez G. et al. 2000) and environmental (Ruiz Corral et al. 2008) classifications. Testing for enrichment of a given variety in a given cluster was performed using Fisher tests with R/stats::fisher.test, under a contingency table formed by partitioning the 1454 accessions by membership of the cluster and assignment to the variety. Results were adjusted using the Holm method with R/stats::p.adjust (α = 0.05).

### Greenhouse evaluation of root anatomy in selected accessions

We selected eight novel native Mexican maize accessions, representative of environmental diversity within the CIMMyT panel for phenotypic validation of RF anatomical predictions. Ten biological replicates of each accession were grown in a greenhouse in State College, PA (40.8028708, -77.8640406) from April to May of 2022. Plants were grown in 2.83 L pots (4 in x 4 in x 14 in, Greenhouse Megastore). The growth media was a mix of silica sand (50%), turface (30%), and field soil (20%) sourced from Rock Springs, PA. Pots were watered to field capacity the night before planting and watered every day after sowing until germination. Once germinated, plants were watered every other day. One week after germination, plants were fertigated with Peters Excel 15 - 5 - 15 Cal Mag Special with Black Iron 200 ppm N recipe and supplemented with an extra 5 ppm Fe (Sprint 330), fed at 1:100 dilution, two times per week until harvest. Greenhouse settings were set at 16 hour days, with a minimum temperature of 21 degrees C and a maximum temperature of 28 degrees C.

Following methods from Burton et al. 2013, 28 days after planting, plants were destructively harvested. Two representative axial roots from nodes two and three were collected. From each axial root, a 4-cm root sample was excised five to nine cm from the most basal portion of the sample. Root samples were stored in 75% ethanol until sectioned by laser ablation tomography (LAT; Strock et al. 2022). In LAT, a sample is moved via an automatic stage towards a 355-nm Avia 7000 pulsed laser and ablated in the focal plane of a camera. A Canon T3i camera with a 53micro lens (MP-E 65 mm) was used to capture images of the root cross-section. Two representative images for each root sample sectioned 1 to 3 cm apart were saved for later image analysis with [RootScan](https://plantscience.psu.edu/research/labs/roots/methods/computer/rootscan). Anatomical phenotypes were averaged for each nodal root of a plant, where each value is an average of two roots from each node and two LAT image sections of each root.

### Estimation of genotypic effects on anatomical traits

To estimate the effect of landrace genotype (race designation) on each measured trait across growth stages (nodes), we used linear mixed models to calculate the best linear unbiased predictions (BLUPs) with the equation:

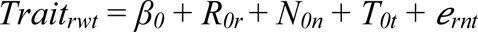

Where for each modeled trait, *Trait_rwt_*, *β_0_* is the overall intercept, *R_0r_* is the random effect of plant race, *N_0n_* is the random effect of node, *T_0t_* is the random effect of the tray the plant was grown in, and *e_rwt_* is the error term. BLUPs were calculated for each measured anatomical trait of a given landrace and extracted using R/lme4::ranef.

### Validation of multivariate fit of trait predictions and observations

We used procrustes analysis to determine concordance between all predicted RF anatomical traits and observed BLUPs. The procrustes analysis relates the overall shape of two sets of multivariate matrices by minimizing the total distance between the two distributions and quantifying how much the relationship between variables in the matrices differ after this alignment (Goodall 1991). The algorithm was implemented using the function R/vegan::procrustes (Oksanen et al. 2022).

### Genome-Wide Association in the Wisconsin Diversity Panel

We used genetic and phenotypic data for 175 inbred maize lines from the expanded Wisconsin Diversity Panel (Schneider et al. 2020). We filtered published SNP data for minor allele frequency >5%, resulting in a total of 370,991 SNPs. Thirteen root anatomical traits were extracted from the published study: MA, MD, MVA, NMV, RXSA, TCA,TSA, AA, X.A, CCFN, CCS, TSA.RXSA, TSA.TCA. The full design and experimental protocol are described in (Schneider et al. 2020). Briefly, maize genotypes were grown in two replicates under well-watered and water-stressed conditions in a randomized complete block design. Plots were irrigated with a center pivot system, and water stress was applied 4 weeks after planting. At anthesis, one representative plant from each plot was excavated from the soil using a standard shovel. Root crowns were soaked and washed to remove soil particles. A representative fourth node root (5-8 cm from the base of the root) was excised and imaged and phenotyped for root anatomical traits using LAT and *RootScan* software. We fitted a linear mixed effect model using R/lme4 (Bates et al. 2007) for the well-watered and water-stressed condition with overall mean as the fixed effect and genotype and block as random effects and extracted BLUPs for genotypes with the R/lme4::ranef function. Broad sense heritability for each root trait was estimated as the genotype variance divided by the sum of genotype variance and error variance from linear mixed effect models. Root trait BLUPs were used to fit linear mixed models using R/GridLMM, a package for fitting linear mixed models with multiple random effects (Runcie & Crawford 2018). We used the function R/GridLMM::GridLMM_GWAS to run the GWA study and set the environmental vector to - 1 or 1 in the model to represent the water-stressed and well-watered treatments. The p-values for the genotype main effect and the genotype by environment interaction effect were calculated using Wald tests. The SNP level P-values were combined into the gene level associations using Multi-marker Analysis of GenoMic Annotation (MAGMA) (de Leeuw et al. 2015). MAGMA uses a multiple regression model to aggregate all SNP information into a gene while accounting for linkage disequilibrium (LD). SNPs were annotated to genes using a 2.5 kilobase window around each gene, resulting in 24,099 genes.

### Environmental GWA

We performed MAGMA to measure the gene-level associations between CIMMyT genotypes and selected environmental variables of their native locations. The first five eigenvectors of the genetic relationship matrix were included in the model to control for population structure. SNPs were annotated to genes using a 2.5 kilobase window around each gene. The final dataset contained 1656 genotypes and 28898 genes for CIMMyT panel accessions.

### Common genes shared between WIDP and CIMMyT

After gene annotation, we obtained 21883 genes that are shared by the WIDP panel and the CIMMyT panel. To evaluate if genes that are highly associated with root anatomical traits also showed associations with our predicted root traits and environmental variables, we extracted and pooled candidate genes from the top 100 genes for all WIDP root traits, predicted root traits, and related environmental variables identified by MAGMA. The final gene list contains WIDP root anatomical genes (576 genotype main effect genes and 542 WIDP genotype x treatment interaction genes), 636 RF predicted anatomical genes, and 1282 environmental genes.

### Minor allele frequency

To understand the relationship between allele variation, environment, and root traits, we extracted the genotypic information of top SNPs of the target genes. We divided maize landraces into PAM clusters, and calculated the mean elevation, the minor allele frequencies (MAF) of the target SNPs, and the mean predicted root traits for each cluster. Pearson correlation was conducted to test the correlations between MAF and elevation, and between MAF and predicted root traits.

## ACKNOWLEDGEMENTS

We acknowledge Kathy Brown and Jonathan Lynch for sharing original root anatomical images and for valuable discussions during the design of this study. We thank Guillaume Lobet (Forschungszentrum Jülich GmbH) and Carlos Ortiz Ramírez (LANGEBIO-CINVESTAV) for providing feedback on the manuscript. We are grateful to the International Maize and Wheat Improvement Center (CIMMYT) for providing seeds. This work was supported by Penn State SNIP2 project *Root system functionality in cereals*. RJHS is funded by USDA Hatch Appropriations under Project #PEN04734 and Accession #1021929. We acknowledge the smallholder farmers and indigenous people whose work and love for their traditions and identity keep maize diversity alive.

## AUTHOR CONTRIBUTIONS

R.J.H.S., C.M.M., and M.L. conceived and designed the analysis. C.M.M. built models used to predict root anatomy; M.L. and R.J.H.S. performed the genetic analysis; A.H. constructed digital anatomies and estimated hydraulic components; C.M.M, M.L., M.P., and R.J.H.S. grew and harvested the pot experiment; C.M.M. obtained anatomical characterizations for novel material using LAT. C.M.M., R.J.H.S., H.S, M.L, and J.L. wrote the paper. All authors read and approved the final version of the manuscript.

## Notes

### Competing Interest Statement

The authors have declared no competing interest.

