## Supplemental Materials for "Evidence that variation in root anatomy contributes to local adaptation in Mexican native maize"

**Supplemental Table 1** Anatomical traits random forest model were constructed for, the Boruta-identified environmental descriptors used in model construction, and the model percent variation explained. Adjusted R-squared is the corrected goodness-of-fit from a linear model between predicted random forest trait values and observed trait values reported in Burton et al., 2013.

| Phenotype | Boruta-identified environmental descriptors | RF model percent variation explained | Adjusted R-squared |
|---|---|---|---|
| TSA.RXSA | prec_3, prec_7, prec_10, prec_11, Ann.Prc, Prc.Wrm.Q | 14.35% | 0.15 |
| TSA.TCA | prec_2, prec_3, prec_7, prec_11, Ann.Prc, Prc.Wet.M, Prc.Wet.Q, Prc.Wrm.Q | 16.69% | 0.18 |
| AA ($mm^2$) | CACO3, BS, PARWin | 33.32% | 0.32 |
| X.A (%) | BS, PARFall | 20.28% | 0.19 |
| X.CCA (%) | org_1m, tmin_4, tmin_10, pet_annual, pet_5, pet_6, pet_9 | 23.12% | 0.21 |
| MA ($mm^2$) | coarse_1m, tmax_2, tmean_3, pet_4 | 23.05% | 0.21 |
| MD (mm) | tmax_2, tmax_3, tmax_4, tmin_3, tmean_3, tmean_5, Ann.Mean.Tmp | 2.12% | 0.05 |
| NMV | tmax_10, tmax_11, tmin_3, tmin_6, tmin_8, tmean_5, tmean_10, tmean_11, Mean.Tmp.Dry.Q, pet_1, pet_12 | 9.46% | 0.16 |
| MVA ($mm^2$) | tmax_1, tmax_2, tmax_3, tmax_4, tmax_11, tmax_12, tmin_3, tmin_4, tmin_5, tmean_2, tmean_3, tmean_4, tmean_5, tmean_11, pet_7, pet_8 | -5.34% | 0.01 |

**Supplemental Table 2** For each CIMMyT PAM cluster assignment, mean elevation, modal assignment of morphological-isozymatic classifications (Ruiz Corral et al. 2008), modal assignment of environmental classifications (Sanchez G. et al. 2000), mean calculated radial conductivity, mean axial, and radial conductance.

| **Cluster** | **Mean elevation (m)** | **Environmental classifications (Ruiz Corral et al. 2008)** | **Morphological-isozymatic classifications (Sanchez G. et al. 2000)** | **Radial conductivity ($k_r$) m MPa$^{-1}$ s$^{-1}$** | **Axial conductance ($K_x$) m$^3$ MPa$^{-1}$ s$^{-1}$** | **Radial conductance ($K_r$) m$^3$ MPa$^{-1}$ s$^{-1}$** |
|---|---|---|---|---|---|---|
| 1 | 502 | Very hot | Tropical dent | $5.92 \times 10^{-5}$ | 7.47 | $2.06 \times 10^{-4}$ |
| 2 | 1912 | Temperate to semi-hot | Highland | $5.87 \times 10^{-5}$ | 4.55 | $2.05 \times 10^{-4}$ |
| 3 | 701 | Very hot | Tropical dent | $5.87 \times 10^{-5}$ | 6.15 | $2.05 \times 10^{-4}$ |
| 4 | 1175 | Temperate to semi-hot | Tropical dent | $5.99 \times 10^{-5}$ | 5.84 | $2.09 \times 10^{-4}$ |
| 5 | 2249 | Temperate to semi-hot | Highland | $5.87 \times 10^{-5}$ | 4.14 | $2.05 \times 10^{-4}$ |
| 6 | 1889 | Temperate to semi-hot | Highland | $6.09 \times 10^{-5}$ | 4.58 | $2.12 \times 10^{-4}$ |
| 7 | 1193 | Temperate to semi-hot | Tropical dent | $5.92 \times 10^{-5}$ | 6.74 | $2.06 \times 10^{-4}$ |

**Supplemental Table 3** Enrichment of Mexican maize varieties in predicted root anatomical PAM clusters.

| Variety | **Environmental classifications (Ruiz Corral et al. 2008)** | **Morphological-isozymatic classifications (Sanchez G. et al. 2000)** | Cluster | Count variety in cluster | Count not variety in cluster | Count variety not in cluster | Count not variety not in cluster | Adj. P-value |
|---|---|---|---|---|---|---|---|---|
| Celaya | 1b: Temperate to Semi-Hot | 3b: Tropical Dent Group | 4 | 33 | 159 | 79 | 1183 | 0.0010 |
| Celaya | 1b: Temperate to Semi-Hot | 3b: Tropical Dent Group | 7 | 24 | 81 | 88 | 1261 | 0.0001 |
| Chalqueño | 1b: Temperate to Semi-Hot | 1a: Highland Cónico Group | 2 | 39 | 260 | 67 | 1088 | 0.0170 |
| Cónico | 1b: Temperate to Semi-Hot | 1a: Highland Cónico Group | 2 | 109 | 190 | 151 | 1004 | 0.0000 |
| Cónico | 1b: Temperate to Semi-Hot | 1a: Highland Cónico Group | 5 | 60 | 59 | 200 | 1135 | 0.0000 |
| Cónico Norteño | 1b: Temperate to Semi-Hot | 1a: Highland Cónico Group | 6 | 43 | 155 | 54 | 1202 | 0.0000 |
| Cristalino de Chihuahua | 1a: Temperate to Semi-Hot, Semi-Dry | 1b: Highland Chihuahua Group | 5 | 9 | 110 | 9 | 1326 | 0.0010 |
| Dzit Bacal | 3: Very Hot | 3a: Mid- to Highland Late Maturity Group | 1 | 13 | 237 | 1 | 1203 | 0.0000 |

| | | | | | | | | |
|---|---|---|---|---|---|---|---|---|
| Nal Tel | 3: Very Hot | 3c: Lowland Short to Medium Maturity Group | 1 | 25 | 225 | 16 | 1188 | 0.0000 |
| Olotillo | 3: Very Hot | 3a: Mid- to Highland Late Maturity Group | 3 | 24 | 267 | 15 | 1148 | 0.0000 |
| Pepitilla | 2c: Semi-Hot to Hot, Semi-Wet | 3c: Lowland Short to Medium Maturity Group | 4 | 18 | 174 | 23 | 1239 | 0.0003 |
| Salvadoreño | N/A | N/A | 1 | 15 | 235 | 1 | 1203 | 0.0000 |
| Tepecintle | 3: Very Hot | 3c: Lowland Short to Medium Maturity Group | 1 | 16 | 234 | 20 | 1184 | 0.0324 |
| Tabloncillo | 2c: Semi-Hot to Hot, Semi-Wet | 2: 8-Row Group | 3 | 25 | 266 | 29 | 1134 | 0.0039 |
| Tuxpeño | 3: Very Hot | 3b: Tropical Dent Group | 1 | 112 | 138 | 184 | 1020 | 0.0000 |
| Vandeño | 3: Very Hot | 3b: Tropical Dent Group | 3 | 19 | 272 | 16 | 1147 | 0.0018 |
| Zapalote Chico | 3: Very Hot | 3c: Lowland Short to Medium Maturity Group | 3 | 11 | 280 | 5 | 1158 | 0.0093 |

**Supplementary Table 4** The gene location and annotation of top overlap genes between WIDP and Prediction GWA.

| GENE | CHR | START | STOP | Predict | WIDP | Gene Name | Gene Symbol |
|---|---|---|---|---|---|---|---|
| **WIDP G and Prediction GWA** | | | | | | | |
| Zm00001d028214 | 1 | 26608063 | 26616524 | MA | TSA.RXSA, TSA.TCA | . | . |
| Zm00001d028831 | 1 | 47987303 | 47994267 | MA | NMV | . | . |
| Zm00001d030005 | 1 | 98582927 | 98597280 | NMV | TSA.RXSA | . | . |
| Zm00001d033829 | 1 | 275155553 | 275162538 | MD | AA | . | . |
| Zm00001d033830 | 1 | 275157506 | 275167900 | MD | AA | . | . |
| Zm00001d002535 | 2 | 14917548 | 14923069 | MA, MD, MVA | TSA.RXSA, TSA.TCA | . | . |
| Zm00001d006052 | 2 | 196573325 | 196584441 | X.A | MA, MD | . | . |
| Zm00001d006053 | 2 | 196580004 | 196586817 | X.A | MA, MD | NAC-transcription factor 24 | nactf24 |
| Zm00001d039901 | 3 | 18642659 | 18653125 | MA | NMV | . | . |
| Zm00001d039902 | 3 | 18643645 | 18649520 | MA, MD | NMV | . | . |
| Zm00001d054106 | 4 | 246772883 | 246783465 | AA, X.A | AA, X.A | Protein disulfide isomerase3 | pdi3 |
| Zm00001d013067 | 5 | 4207061 | 4213378 | X.A | X.A | . | . |
| Zm00001d015376 | 5 | 87049440 | 87058148 | AA, X.A | MA, MD | Phosphoglycerate kinase3 | pgk3 |
| Zm00001d015397 | 5 | 88303760 | 88309464 | MVA, NMV | MA, MD | VQ motif-transcription factor29 | vq29 |

| | | | | | | | |
|---|---|---|---|---|---|---|---|
| Zm00001d016842 | 5 | 178207500 | 178214684 | TSA.RXSA, TSA.TCA | NMV | . | . |
| Zm00001d036241 | 6 | 78621119 | 78628756 | MD | NMV | . | . |
| Zm00001d012089 | 8 | 168026879 | 168032905 | MVA, NMV | X.A | . | . |
| Zm00001d012090 | 8 | 168029331 | 168034798 | MVA, NMV | X.A | . | . |
| **WIDP GxE and Prediction GWA** | | | | | | | |
| Zm00001d028211 | 1 | 26522710 | 26532325 | AA, X.A | MA, MD, MVA | . | . |
| Zm00001d001833 | 2 | 1623622 | 1634808 | MD | X.A | . | . |
| Zm00001d003660 | 2 | 52983973 | 52990152 | NMV | X.A | . | . |
| Zm00001d003661 | 2 | 52985264 | 52992526 | NMV | X.A | . | . |
| Zm00001d039542 | 3 | 7733246 | 7739763 | MD, MVA, NMV | MVA | High PhosphatidylCholine1 | hpc1 |
| Zm00001d044130 | 3 | 220026982 | 220032889 | TSA.RXSA, TSA.TCA | MA, MD, MVA | . | . |
| Zm00001d049894 | 4 | 51234927 | 51283694 | MD, MVA | MD | . | . |
| Zm00001d053932 | 4 | 243765590 | 243773531 | NMV | AA, X.A | . | . |
| Zm00001d021059 | 7 | 141817446 | 141823675 | TSA.RXSA, TSA.TCA | MA, MD, MVA | Stearoyl-acyl-carrier-protein desaturase8 | sacd8 |
| Zm00001d021060 | 7 | 141819400 | 141826721 | TSA.RXSA, TSA.TCA | MA, MVA | . | . |
| Zm00001d008900 | 8 | 24521219 | 24526917 | NMV | MA, MD | . | . |

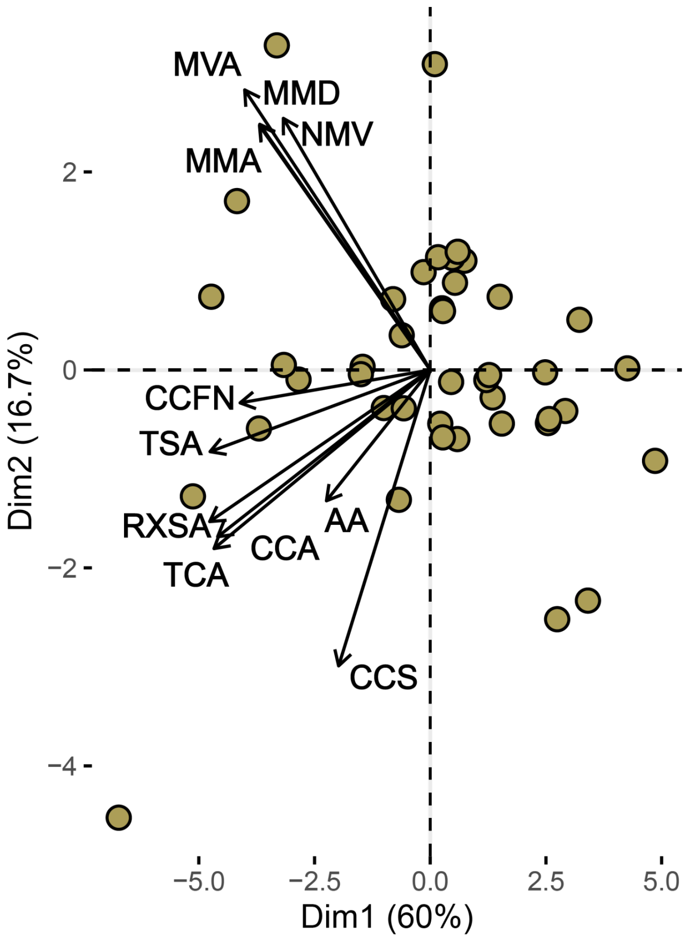


**Supplemental Figure 1** PC1 vs PC2 loadings of main root anatomical traits for accessions from the Burton panel. Dimension 1 primarily delineates smaller root cross-sections from smaller root cross-sections.

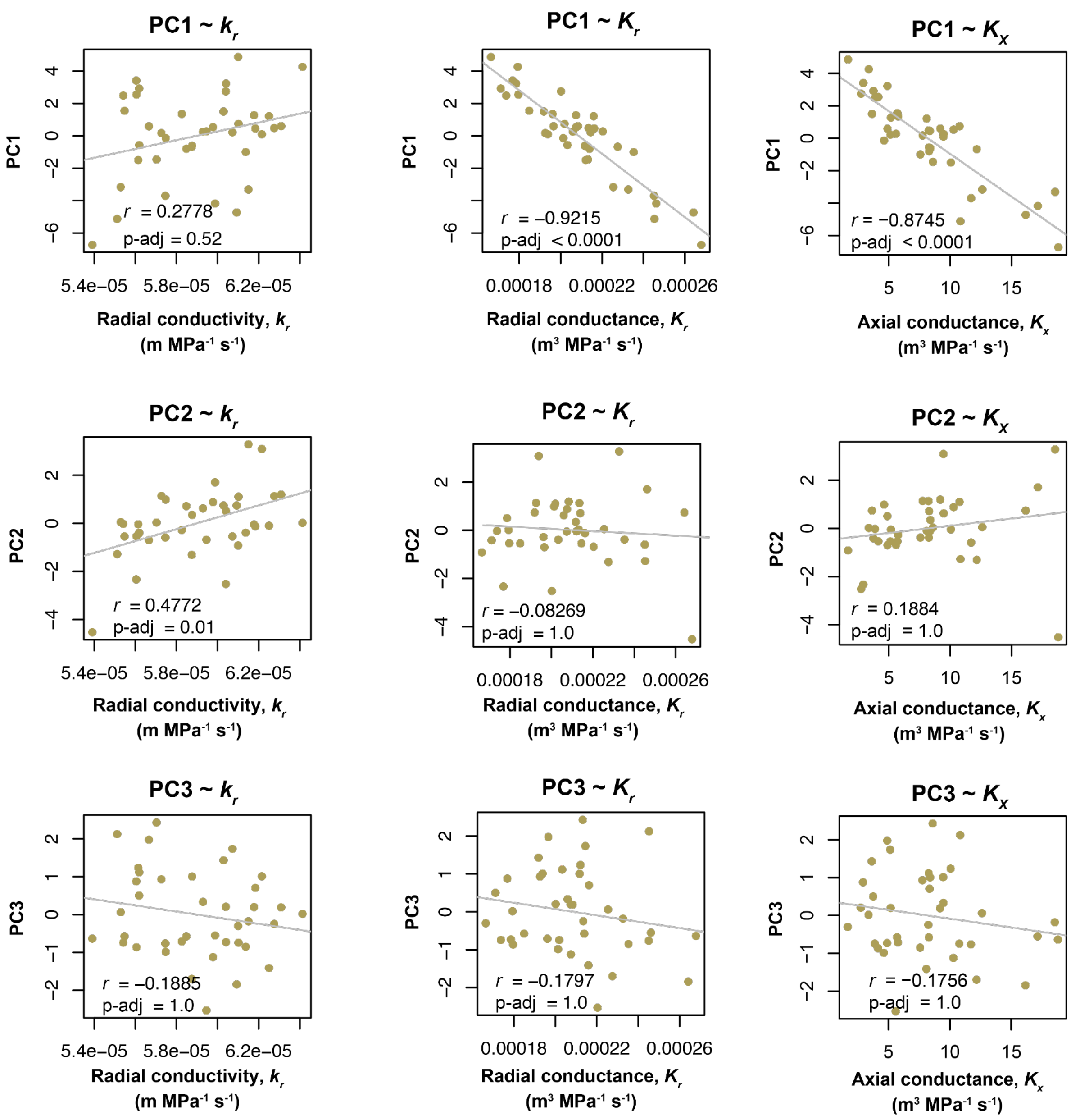


**Supplemental Figure 2** Loadings of Burton panel anatomical PC1, PC2, and PC3 versus calculated radial conductivity ($k_r$), radial conductance ($K_r$,), and axial conductance ($K_x$) of the same accessions. P-values for each PC dimension are adjusted using the Holm-Bonferroni method.

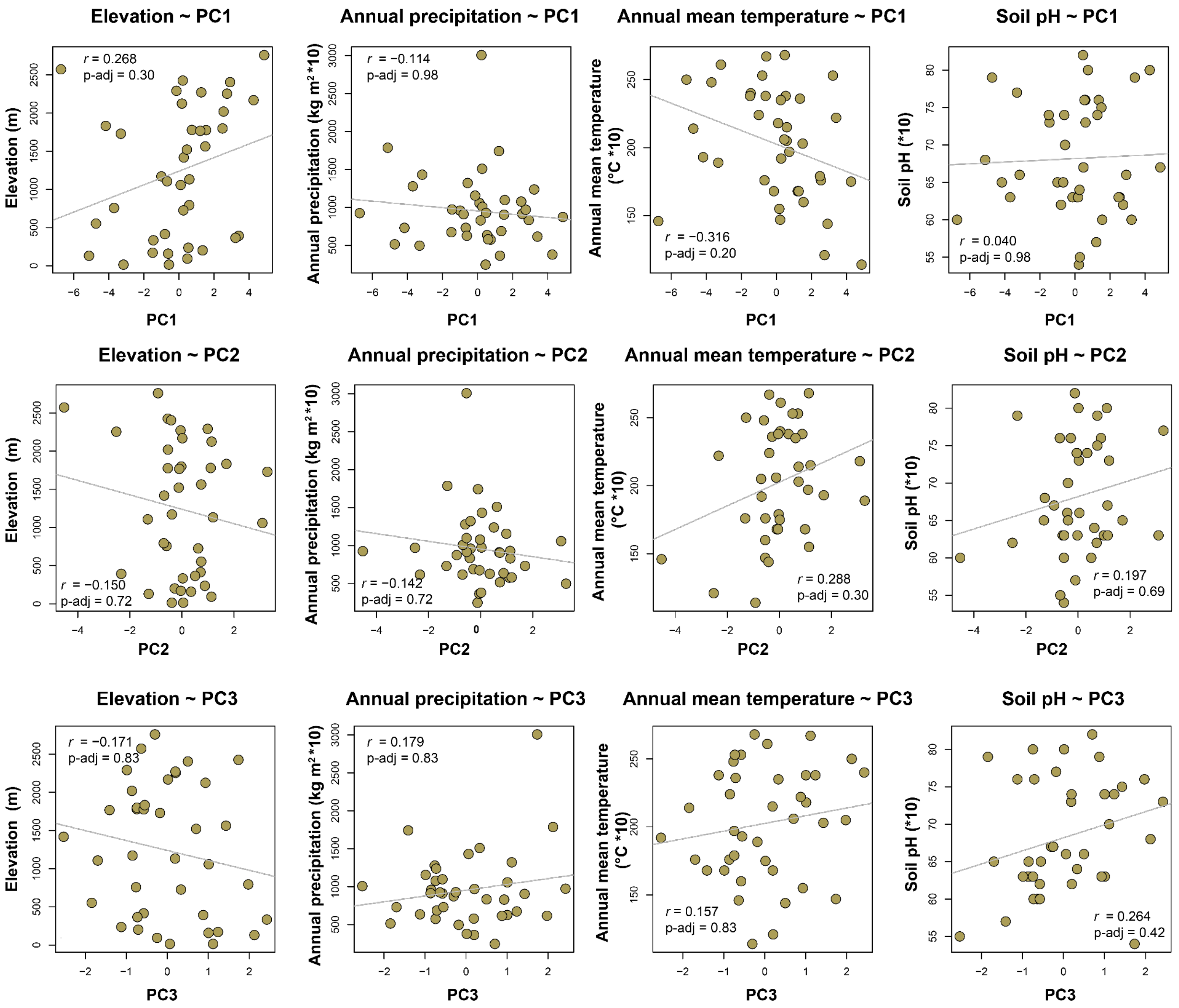


**Supplemental Figure 3** Four core environmental aspects of the Burton panel point of origin (elevation, annual precipitation, mean temperature, and soil pH) versus loadings of Burton panel anatomical PC1, PC2, of the same accessions. P-values for each PC dimension are adjusted using the Holm-Bonferroni method.

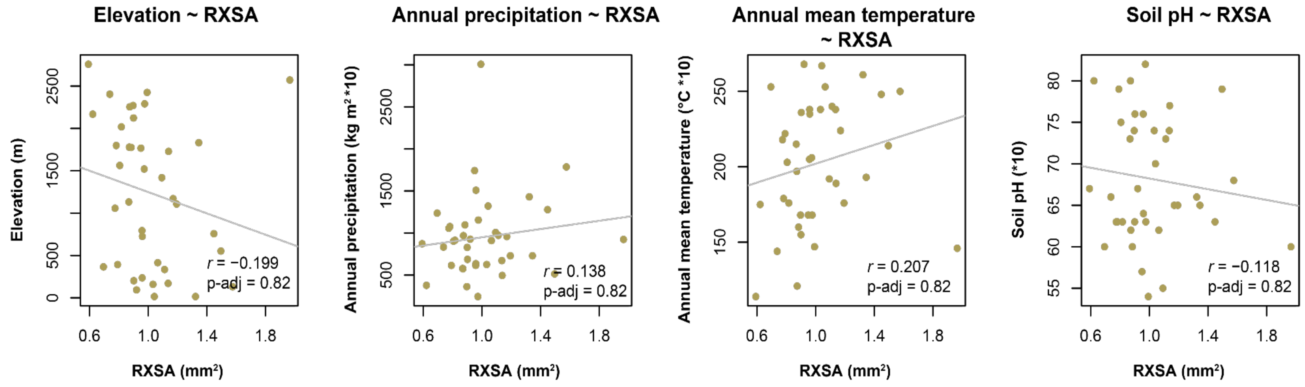


**Supplemental Figure 4** Four core environmental aspects of the Burton panel point of origin (elevation, annual precipitation, mean temperature, and soil pH) versus root cross section of Burton panel accessions. P-values are adjusted using the Holm-Bonferroni method.

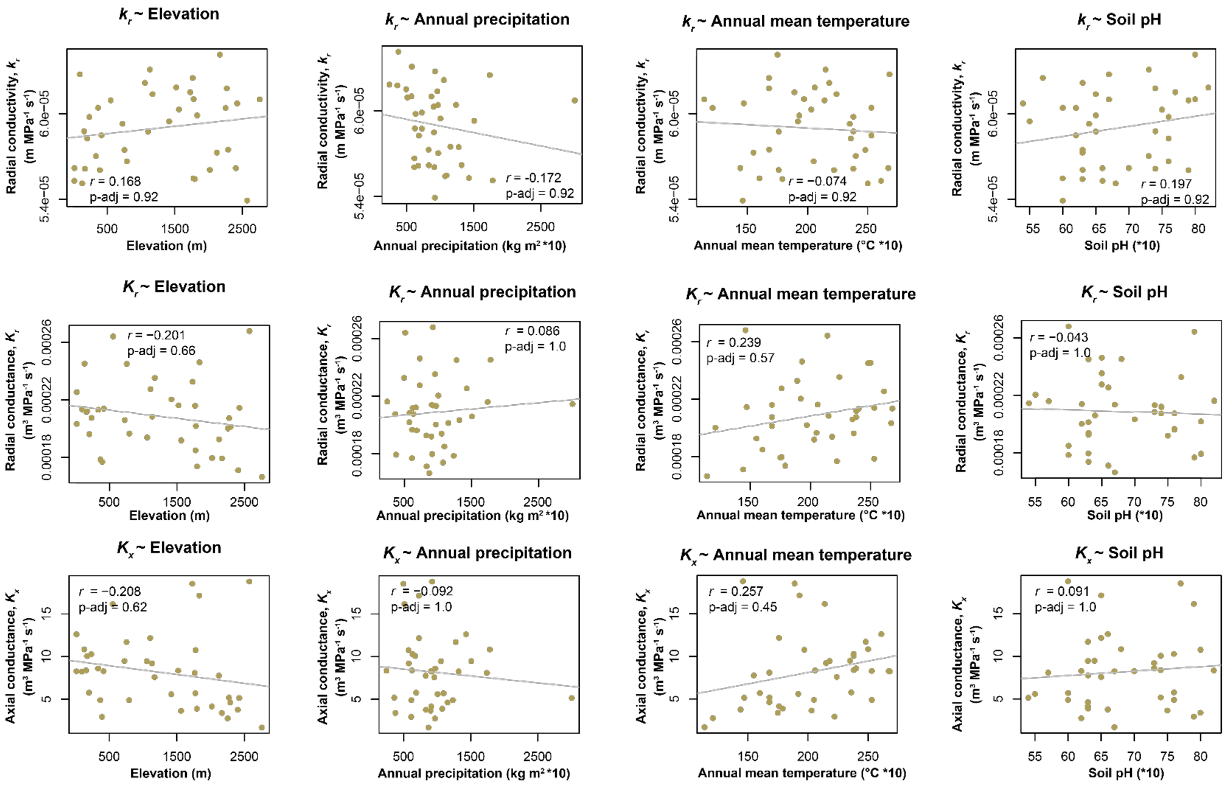

**Supplemental Figure 5** Calculated radial conductivity (*$k_r$*), radial conductance (*$K_r$*,), and axial conductance (*$K_x$*) versus four core environmental aspects of the Burton panel point of origin (elevation, annual precipitation, mean temperature, and soil pH). P-values for each hydraulic scenario are adjusted using the Holm-Bonferroni method.

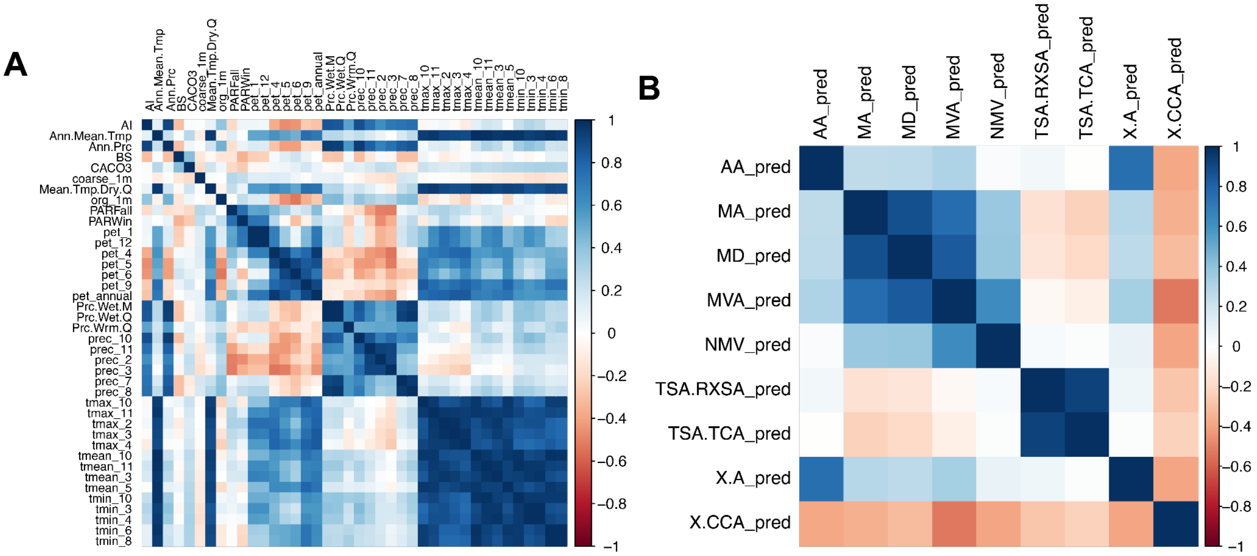


**Supplemental Figure 6** Correlation plots of A) environmental descriptors used across all RF models and B) RF predicted anatomical traits of the CIMMyT panel. Blue colors indicate positive and red colors indicate negative correlations between variables.

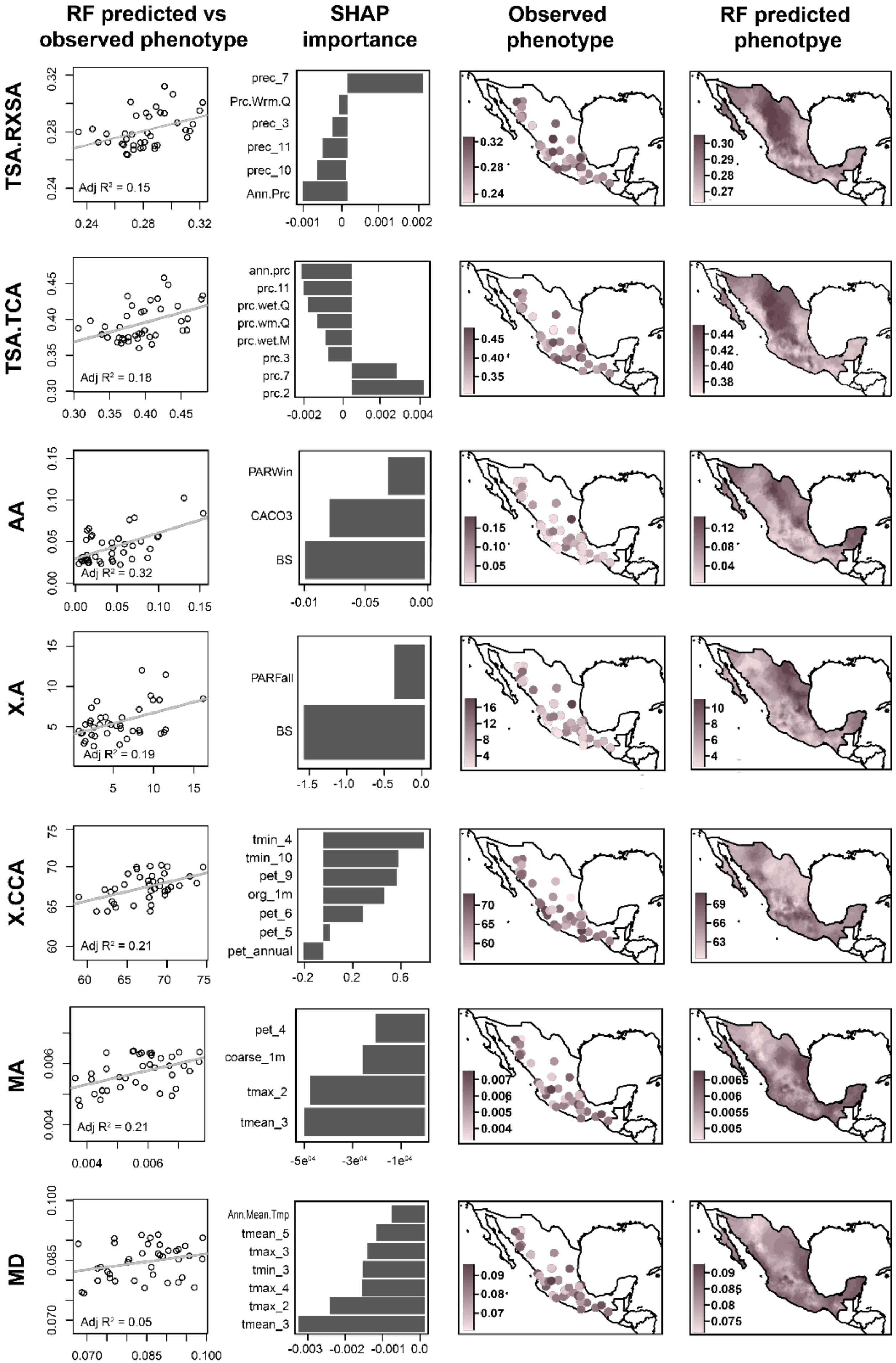

RF predicted vs observed phenotype
SHAP importance
Observed phenotype
RF predicted phenotpye
TSA.RXSA
prec_7
Prc.Wrm.Q
prec_3
prec_11
prec_10
Ann.Prc
Adj R² = 0.15
TSA.TCA
ann.prc
prc.11
prc.wet.Q
prc.wm.Q
prc.wet.M
prc.3
prc.7
prc.2
Adj R² = 0.18
AA
PARWin
CACO3
BS
Adj R² = 0.32
X.A
PARFall
BS
Adj R² = 0.19
X.CCA
tmin_4
tmin_10
pet_9
org_1m
pet_6
pet_5
pet_annual
Adj R² = 0.21
MA
pet_4
coarse_1m
tmax_2
tmean_3
Adj R² = 0.21
MD
Ann.Mean.Tmp
tmean_5
tmax_3
tmin_3
tmax_4
tmax_2
tmean_3
Adj R² = 0.05

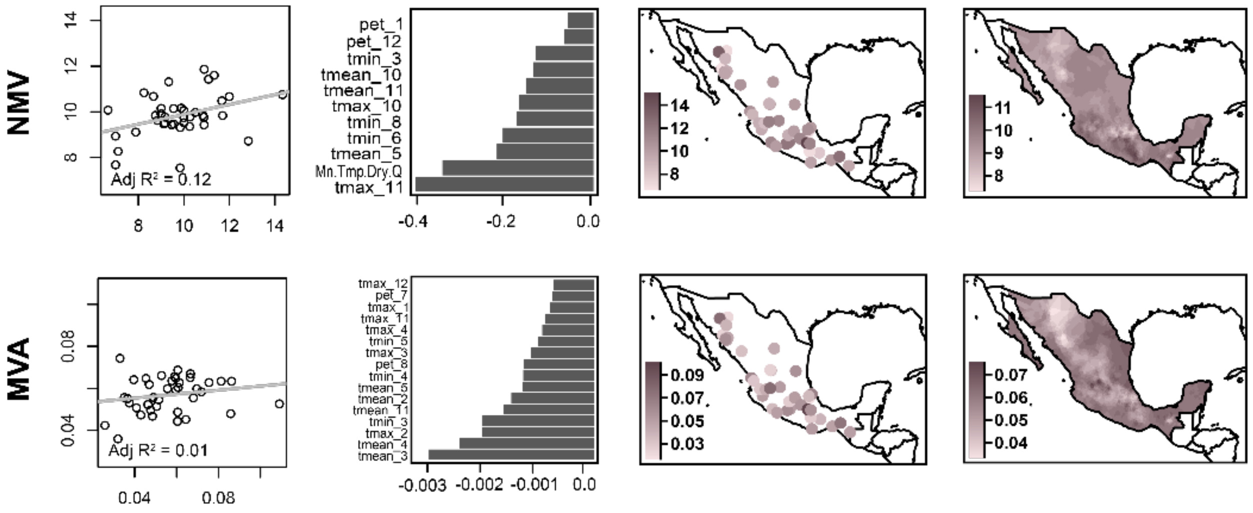


**Supplemental Figure 7** RF predicted vs observed values for individuals used in model training and validation. Line is the coefficient of determination for all plotted points. SHAP values (SHapley Additive exPlanations) of environmental descriptors used in model training, describing the contribution and direction of each environmental descriptor used in a RF model. Burton panel accession point of origin colored by reported trait observations used to train RF models. Smoothed trait predictions as a function of the environment from constructed RF models.

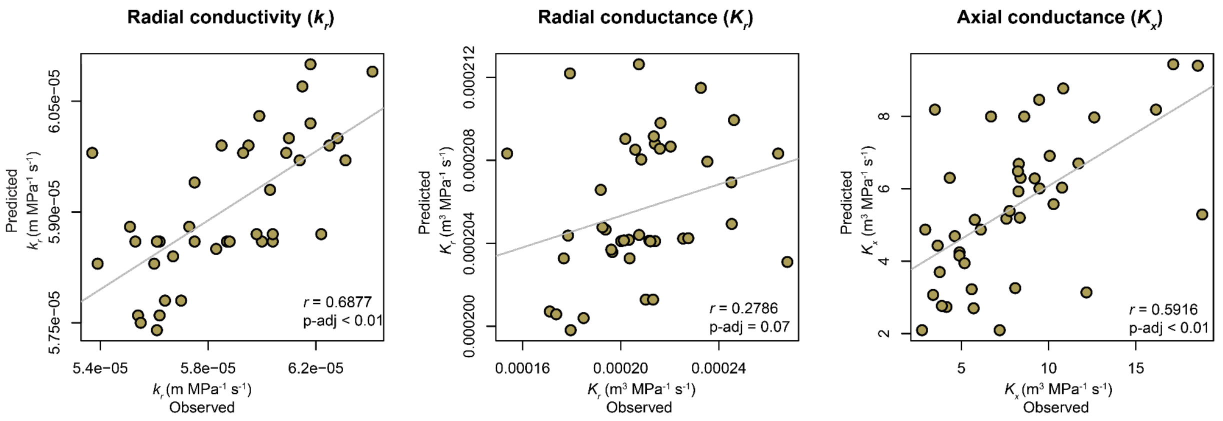


**Supplemental Figure 8** Observed vs predicted radial conductivity ($k_r$), radial conductance ($K_r$), and axial conductance ($K_x$) for the Burton panel accessions. Radial conductivity is the same as shown in Figure 2D. P-values are adjusted using the Holm-Bonferroni method.

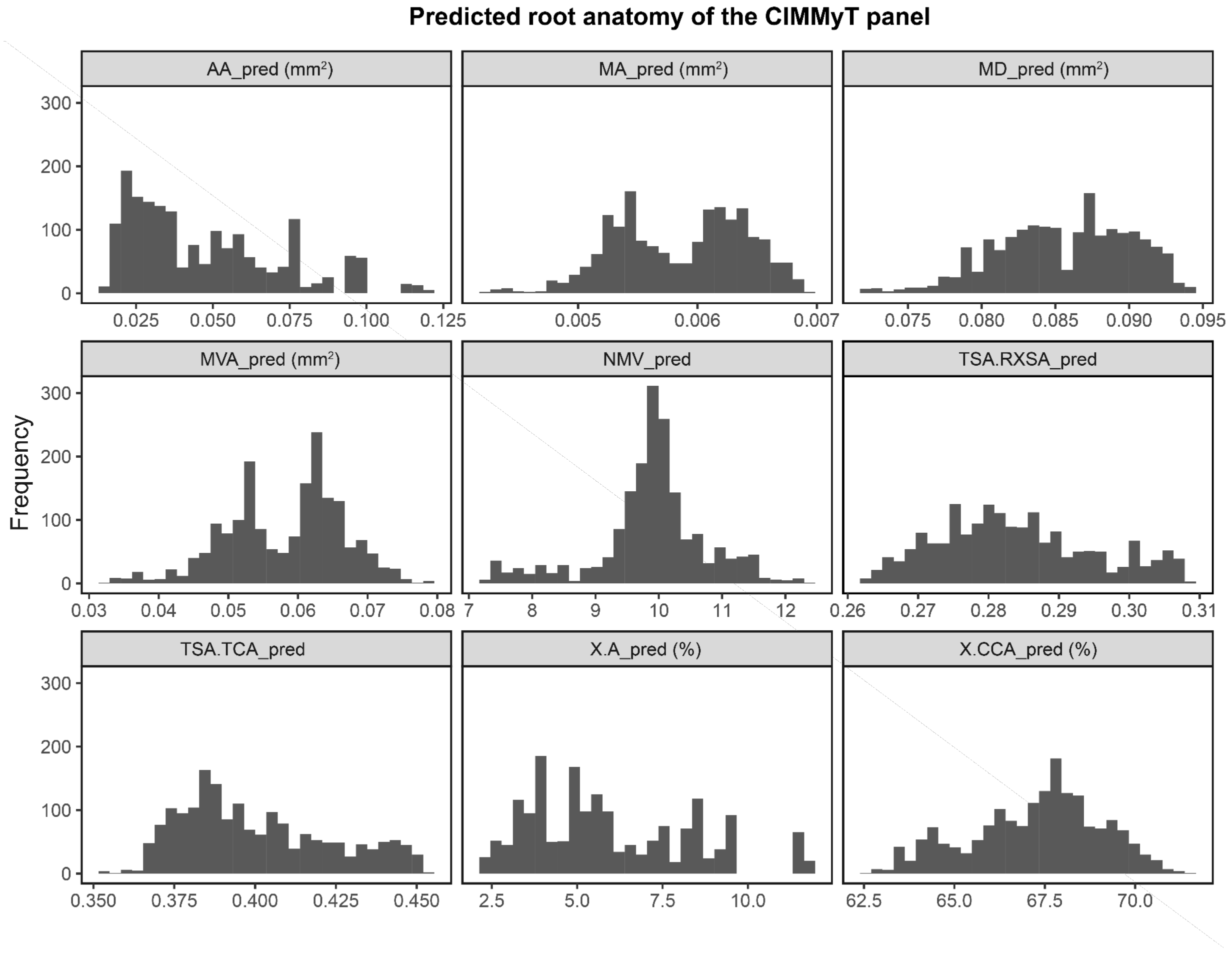


**Supplemental Figure 9** Histograms of RF anatomical trait predictions for the CIMMyT panel.

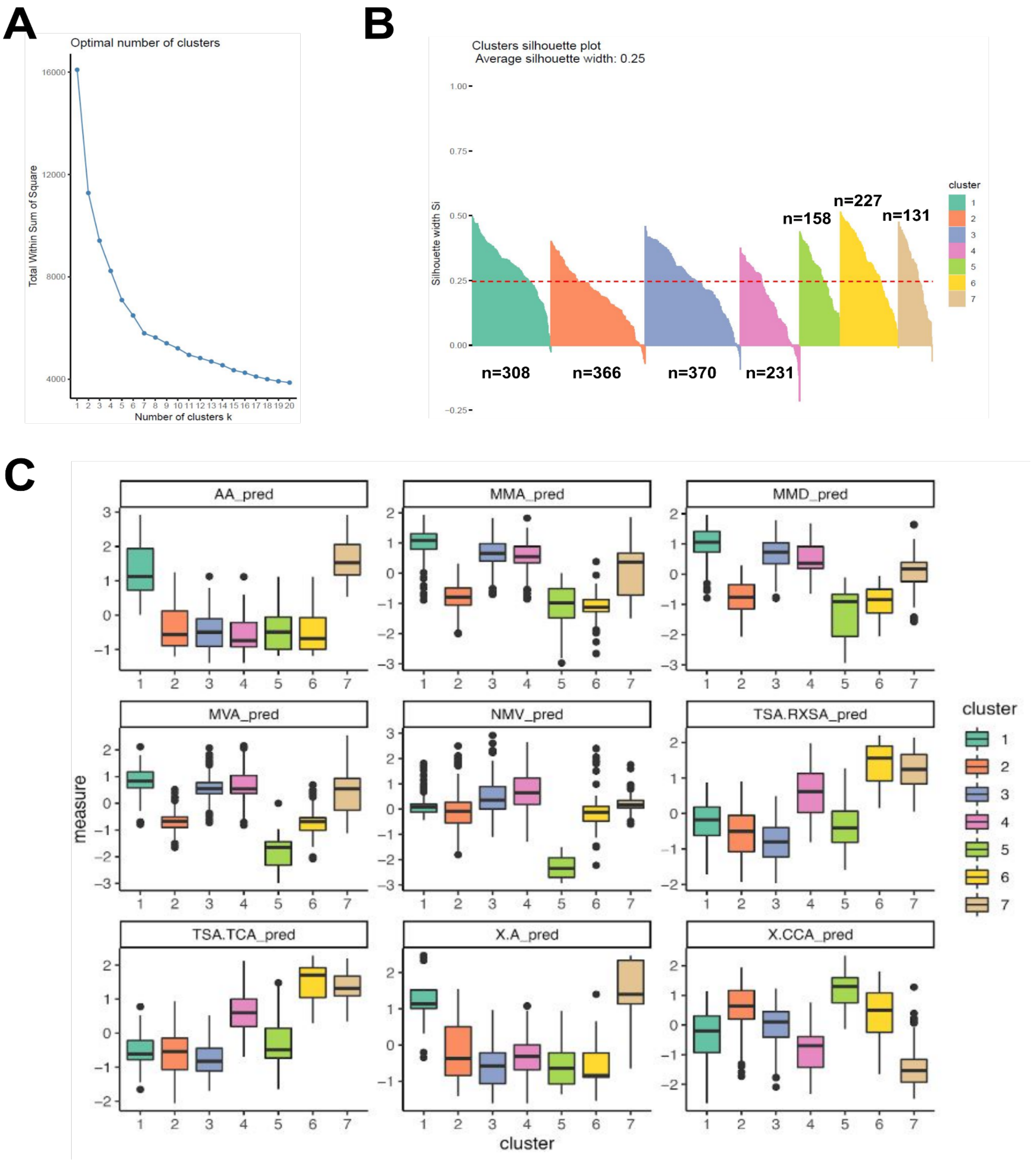


**Supplemental Figure 10** A) Within cluster sums of squares for predicted anatomy of the CIMMyT panel. Used to determine the optimal number of clusters for partitioning against means (PAM). B) Cluster silhouettes summarizing the degree of separation between the clusters. The number of accessions in each cluster is shown. C) Boxplots by cluster for all RF predicted CIMMyT traits.

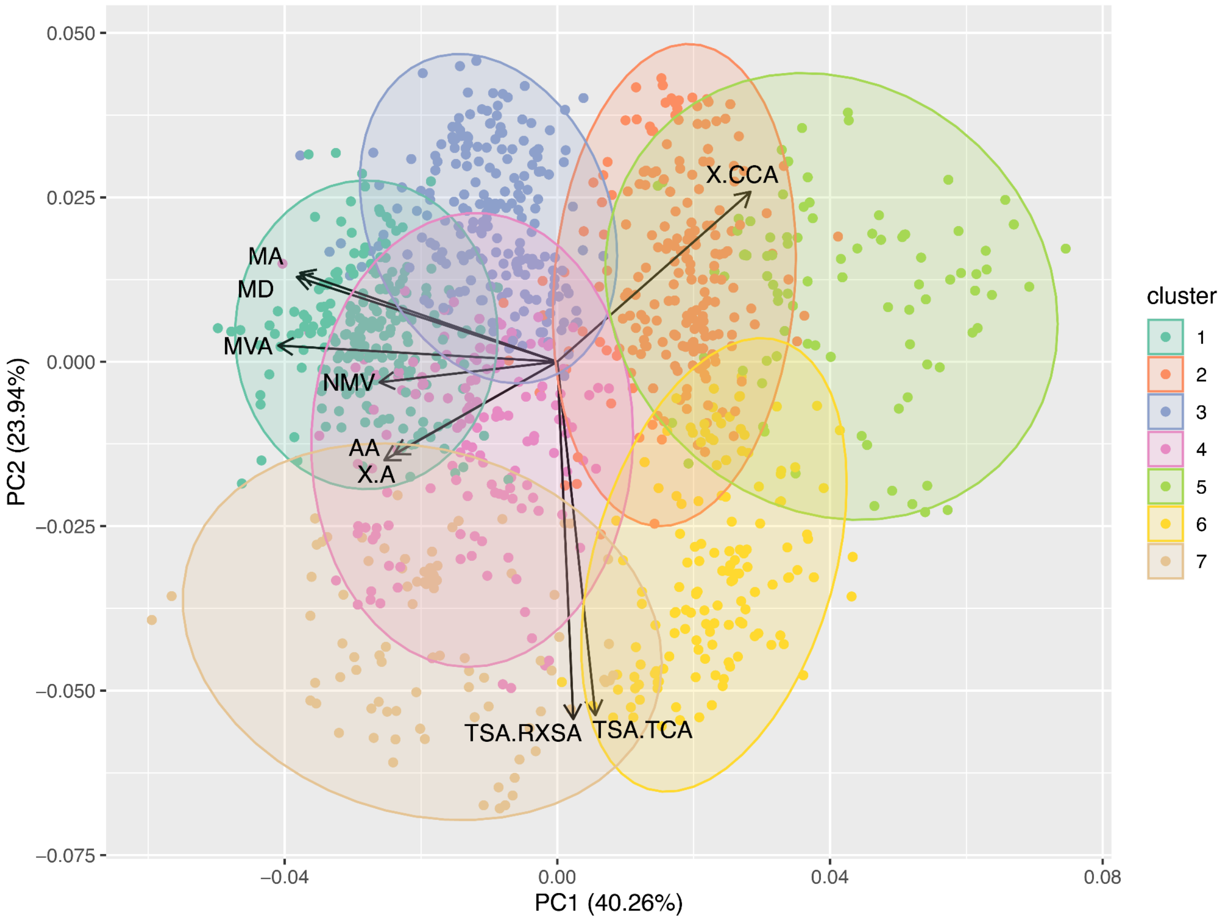


**Supplemental Figure 11** PC plot of RF predicted traits for individuals of the CIMMyT panel used in PAM clustering. Clusters are defined by differences in cortical and metaxylem traits.

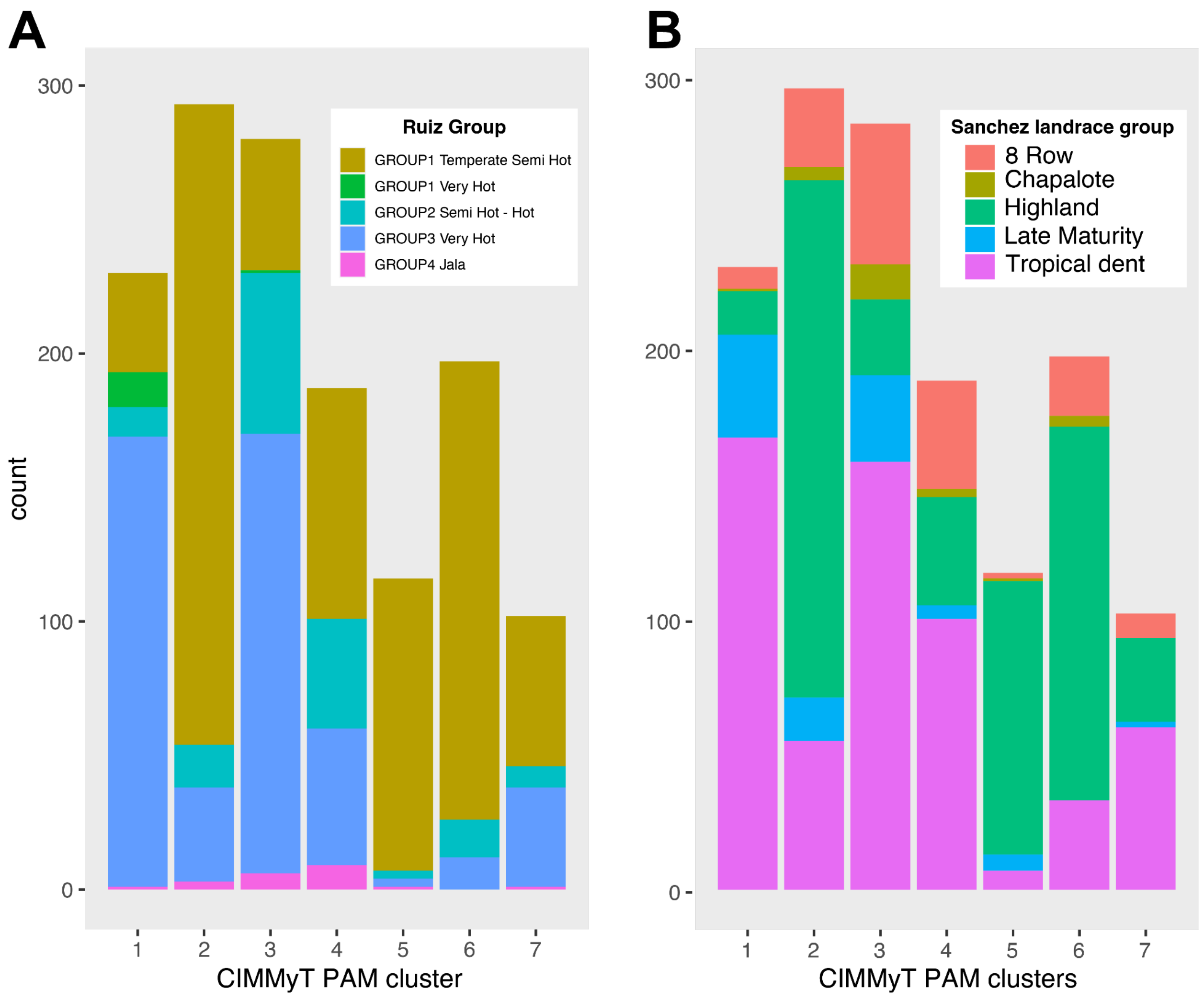


**Supplemental Figure 12** Distribution of CIMMyT PAM cluster individuals using landrace designation, grouped by previous A) environmental (Ruiz Corral et al. 2008) classifications and B) morphological-isozymatic (Sanchez G. et al. 2000). For each cluster, the primary morphological-isozymatic and environmental classification is reported in Supplemental Table 3.

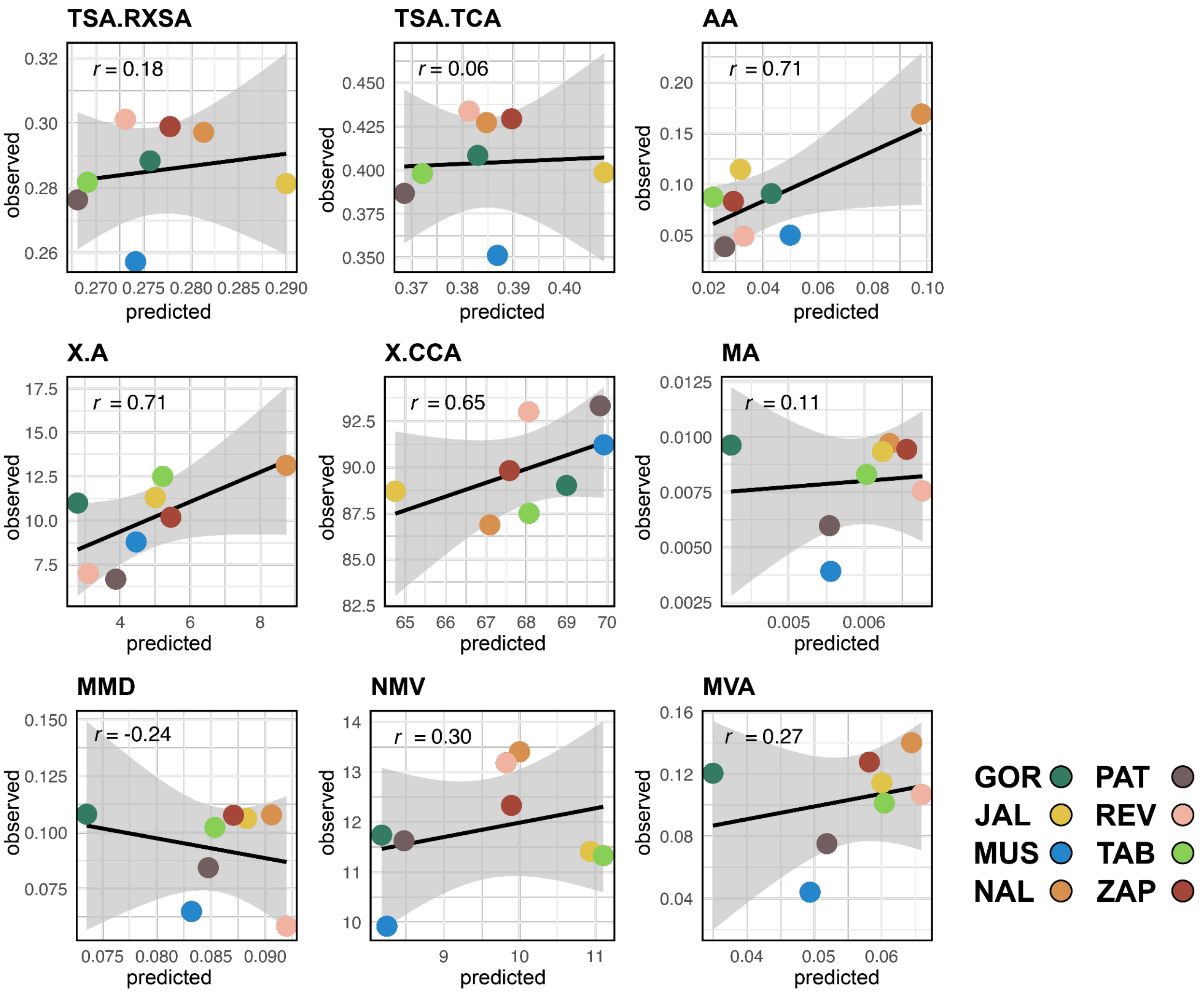


**Supplemental Figure 13** Validation lines observed BLUPs of nodal root two and three from greenhouse grown plants vs RF trait predictions.

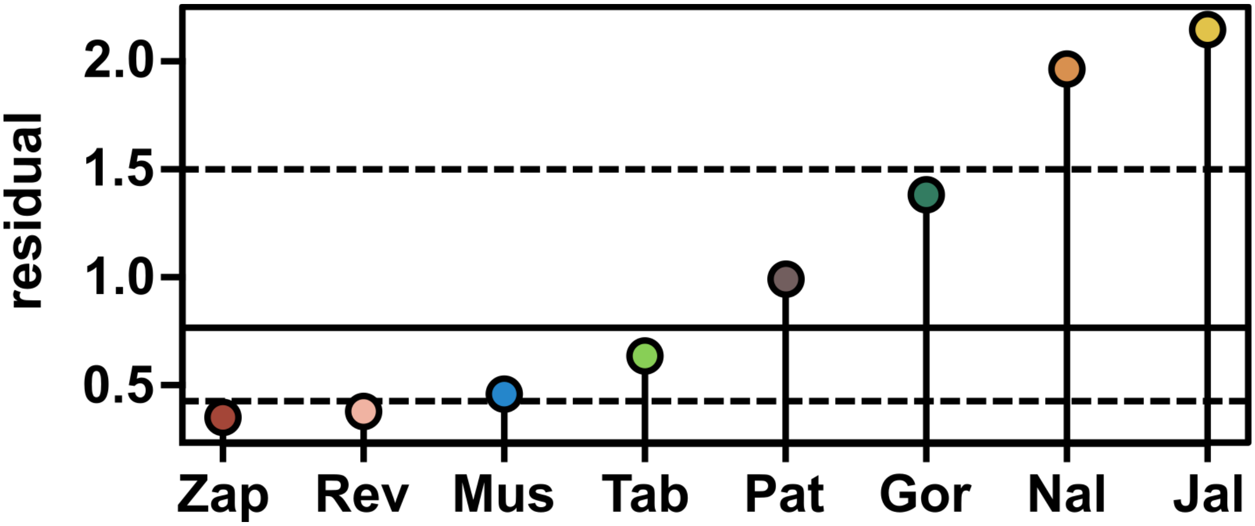


**Supplemental Figure 14** Procrustes errors of comparing principal component analysis (PCA) of observed anatomical triats of novel material grown in this study and RF anaomtical trait predictions (ordered according to the residual scores). Dashed and solid lines are the first, second, and third quartiles.

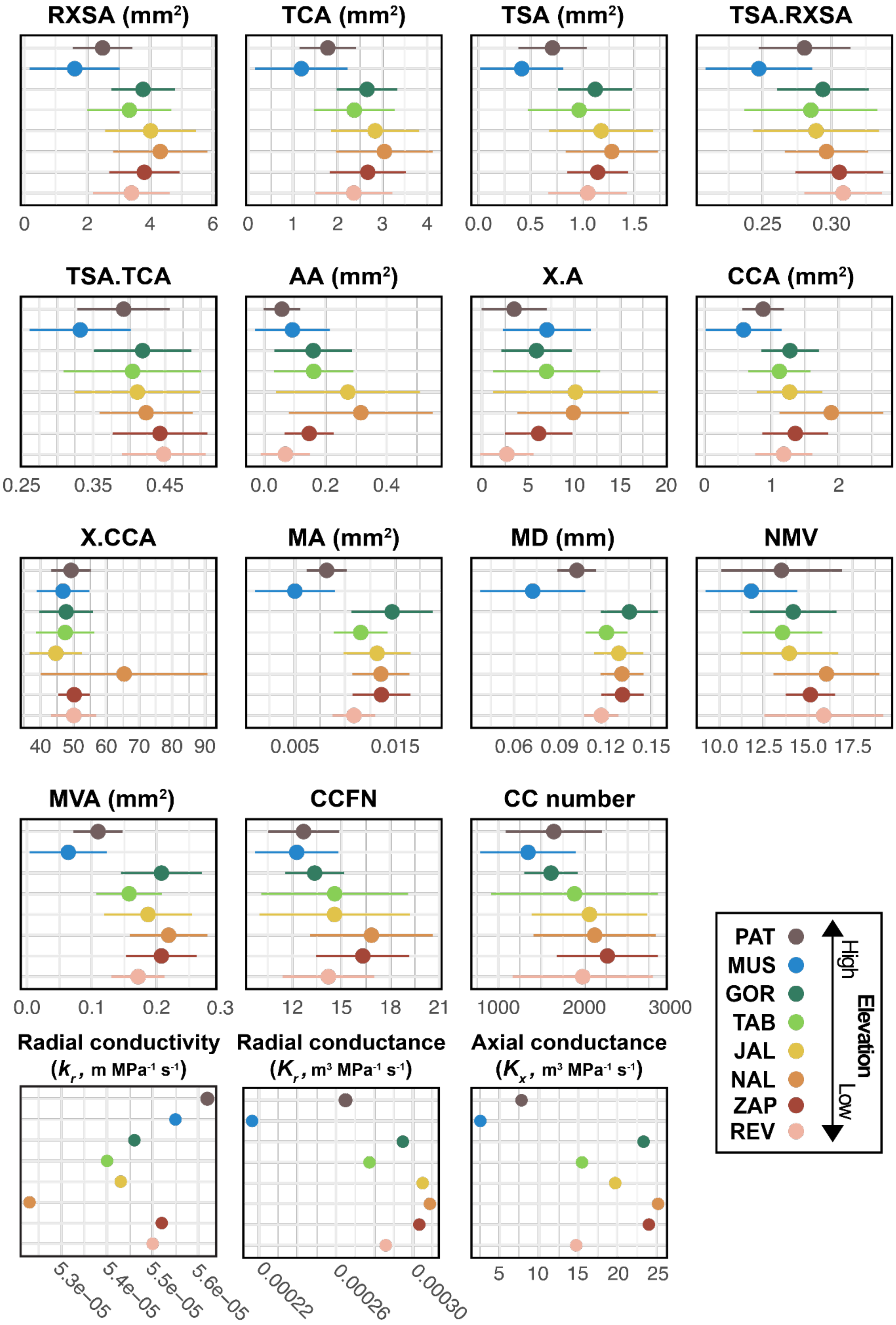

RXSA (mm²)
TCA (mm²)
TSA (mm²)
TSA.RXSA
TSA.TCA
AA (mm²)
X.A
CCA (mm²)
X.CCA
MA (mm²)
MD (mm)
NMV
MVA (mm²)
CCFN
CC number
Radial conductivity ($k_r$, m MPa⁻¹ s⁻¹)
Radial conductance ($K_r$, m³ MPa⁻¹ s⁻¹)
Axial conductance ($K_x$, m³ MPa⁻¹ s⁻¹)
PAT
MUS
GOR
TAB
JAL
NAL
ZAP
REV
High
Elevation
Low

**Supplemental Figure 15** Observed whorl three anatomical traits of novel material grown in this study. Genotypes ordered by elevation. Each line represents the maximum and minimum trait value observed for a genotype with the circle as the genotype mean. Conductance and conductivity values are represented as the mean value for the genotype modeled by MECHA.

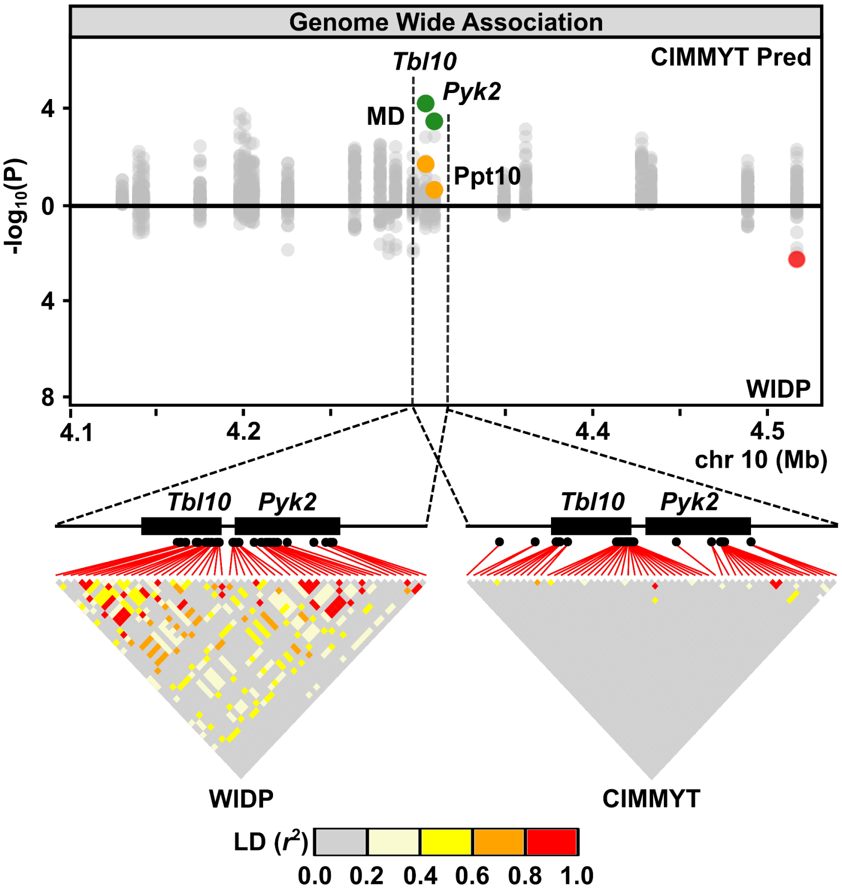


**Supplemental Figure 16:** Miami plot showing GWA support (-log$_{10}$P) for association with root anatomy for genes on a region of chromosome 10. Points above the x-axis show support for predicted phenotypes in the CIMMyT panel; points below the x-axis show support for observed phenotypes for the WIDP panel. Two genes (*Tbl10* and *Pyk2*) in the highlighted region show strong support for *mean metaxylem vessel diameter* (MD) in the CIMMyT panel, but no signal for the WIDP panel. Image below the Miami plot shows *Tbl10* and *Pyk2* genes in genomic context (CDS as filled boxes), SNP position (filled circles) and pairwise linkage disequilibrium (LD) for the CIMMyT and WIDP marker.
